# Neutrophil Extracellular Trap gene expression signatures identify prognostic and targetable signaling axes for inhibiting metastasis of pancreatic tumours

**DOI:** 10.1101/2024.11.15.622336

**Authors:** Paul C. McDonald, James T Topham, Shannon Awrey, Hossein Tavakoli, Rebekah Carroll, Wells S. Brown, Zachary J. Gerbec, Steve E. Kalloger, Joanna M. Karasinska, Patricia Tang, Rachel Goodwin, Steven J. M. Jones, Janessa Laskin, Marco A Marra, Gregg B. Morin, Daniel J. Renouf, David F. Schaeffer, Shoukat Dedhar

## Abstract

Tumour associated neutrophils (TANs) can promote metastasis through the interaction of Neutrophil Extracellular Traps (NETs) with tumour cells. Here, we examined the contribution of NETs in the progression of Pancreatic Ductal Adenocarcinoma (PDAC), which is characterized by high metastatic propensity. We carried out consensus clustering and pathway enrichment analysis of NET-related genes in an integrated cohort of 369 resectable and metastatic PDAC patient tumour samples, and compiled two gene expression signatures comprising of either, integrin-actin cytoskeleton, and Epithelial to Mesenchymal Transition (EMT) signaling, or cell death signaling, which identified patients with very poor to better overall survival, respectively. Tumour Infiltrating neutrophils and NETs associate with ITGB1, CCDC25 and ILK, within clinical and experimental PDAC tumours. Functionally, exposure of PDAC cells to NETs identified a cytoskeletal dynamic-associated CCDC25/ITGB1/ILK signaling complex which stimulates EMT and migration/ invasion. Furthermore, NETosis-driven experimental metastasis of PDAC cells is significantly inhibited by *ILK* knock down. Our data identify novel NET-related gene expression signatures for PDAC patient stratification, and reveal targetable signaling axes to prevent and treat disease progression and metastasis.

---

Pancreatic Ductal Adenocarcinoma (PDAC) is a difficult-to-treat cancer punctuated by early metastasis and a dismal prognosis^1–3^. Chronic pancreatitis is a risk factor for the development of PDAC^4,5^, and tumours and metastases are characterized by an immunosuppressive, fibro-inflammatory tumour microenvironment (TME), underscoring the importance of inflammation as a driver of PDAC progression^6,7^. Neutrophils, vital cellular effectors of inflammation^8^, have emerged as critical pro-tumourigenic and metastasis mediators^9–11^ and neutrophils contribute to tumour progression and metastasis, including “awakening” dormant circulating cancer cells ^12–16^.

Neutrophils can undergo NETosis in response to cytokine or chemokine mediated inflammatory stimuli to form neutrophil extracellular traps (NETs), which are structures composed of decondensed, modified genomic DNA, histones and proteases^17,18^. NETs have been shown to interact with tumour cells within primary tumour, within the vasculature, or at the metastatic site^13–15,19^. The NET-tumour interaction promotes metastatic colonization by stimulating both invasion as well as growth within the metastatic site^14,15,20,21^.

While studies in PDAC have shown that NETs are present both in primary tumours and in metastases^22,23^, the molecular details of the interactions between NETs and PDAC cells are poorly understood, and the contribution of neutrophils and NETs to PDAC progression and metastasis remains unclear.

Here we set out to determine, using unbiased genomic and proteomic analysis of resectable and metastatic PDAC patient cohort data, the role of NETosis and NETs in PDAC patient outcome. We have identified novel prognostic NET gene expression signatures that reveal NET-associated ITGB1/CCDC25/ILK mediated cytoskeletal dynamics and EMT pathways as poor outcome indicators for PDAC patients. Our data demonstrate a novel association of CCDC25 with ITGB1 and ILK in the context of NET-mediated cytoskeletal reorganization and cell migration/invasion. Furthermore, we demonstrate that patient and experimental PDAC tumours and metastases harbour significant amounts of intra tumoural neutrophils with associated NETs. Functional exploration of whether the NET signature-related ITGB1/CCDC25/ILK/ and EMT axis plays a role in the response of PDAC cells to NETs reveals that this axis promotes NET-induced PDAC cell invasion and metastasis, suggesting novel prognostic and therapeutic approaches for the prevention and inhibition of PDAC metastasis.

## Results

### Neutrophil extracellular trap (NET) gene signatures identify PDAC patient outcomes

Pan-cancer analyses across 14 solid cancer types have shown that, amongst all tumour-infiltrating leukocyte populations, intratumoural neutrophils are most robustly associated with poor prognosis^24^ and a high circulating neutrophil-to-lymphocyte ratio (NLR) is also correlated with shorter survival^10,11^, signaling the importance of tumour-infiltrating neutrophils in cancer progression. To determine whether a higher NLR is relevant to the outcome of PDAC patients in an advanced disease setting, we examined the baseline circulating NLR in the blood samples of a cohort of 67 patients presenting with metastatic PDAC. We observed that a higher baseline NLR is associated with significantly shorter overall patient survival (Fig. 1A), establishing a clinical link between neutrophils and metastatic PDAC.

**Figure 1.**
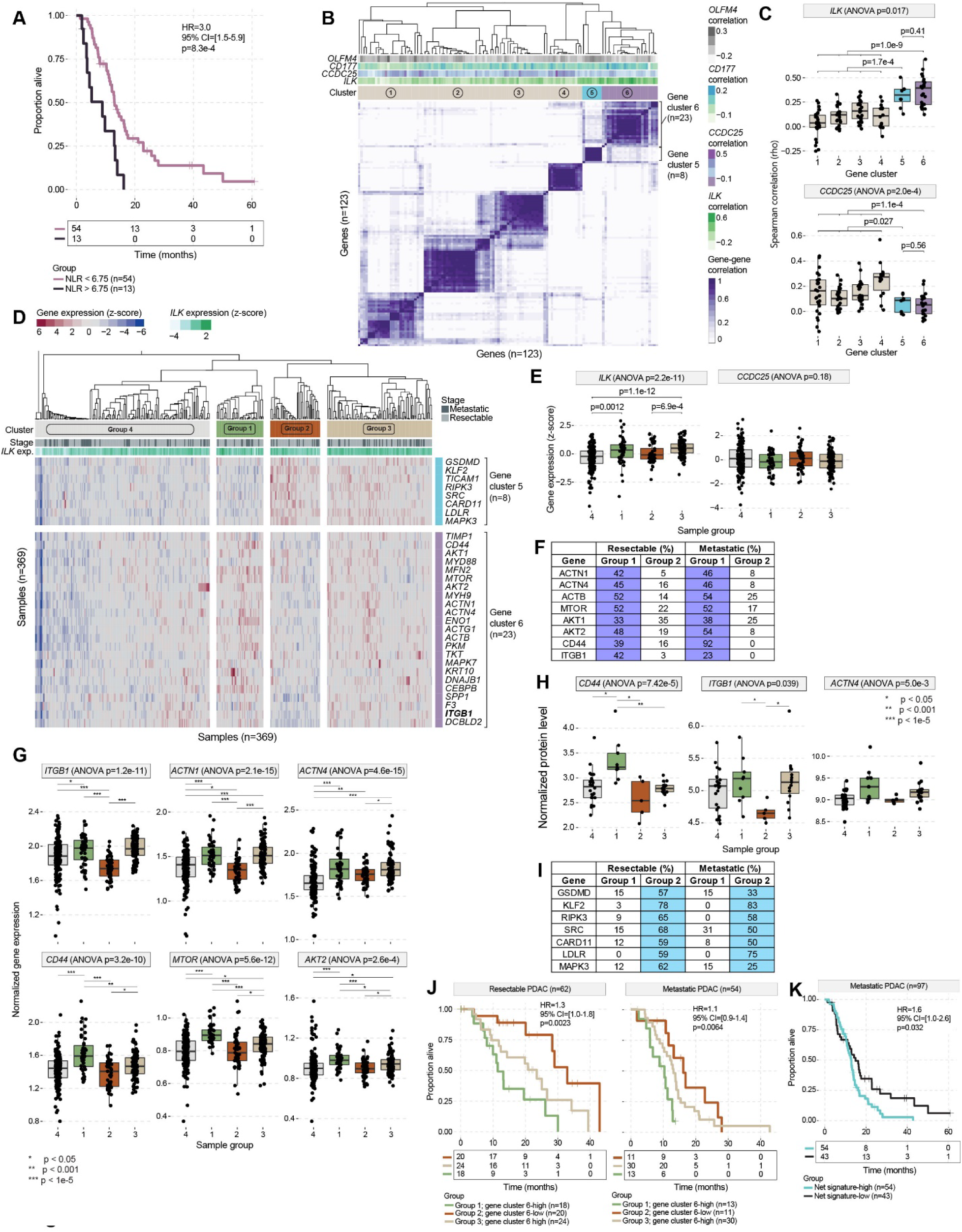
A neutrophil extracellular trap (NET) gene signature correlating with high ILK expression identifies a group of PDAC patients with poor prognosis. **(A)** Kaplan Meier survival analysis showing that a higher baseline neutrophil-to-lymphocyte ratio (NLR) is associated with shorter overall survival in patients with metastatic PDAC (n = 67). **(B)** Heat map showing the optimal consensus clustering solution for NET-related genes in human PDAC samples (resectable and metastatic combined, n = 369). Tracks above the clusters show the Spearman correlation (rho) of the indicated genes versus each gene in the heat map. **(C)** Box plots showing the relationship between expression of the indicated genes and each gene cluster identified in panel B. **(D)** Heat map showing clustering of human PDAC samples (n = 369) according to expression (z-score) of genes comprising a NET signature defined by consensus clustering in Panel B. **(E)** Box plots showing levels of gene expression (z-score) for the indicated genes in each of the sample clusters identified in Panel D. **(F)** Table showing the proportion of samples for patient groups 1 and 2 that have upregulation of the indicated genes from gene cluster 6. **(G, H)** Box plots showing (G) transcript levels and (H) protein levels for a subset of genes across the different patient groups identified in Panel D. **(I)** Table showing the proportion of samples for patient groups 1 and 2 that have upregulation of the indicated genes from gene cluster 5. **(J)** Kaplan-Meier survival analysis of patients with resectable (left) and metastatic (right) PDAC stratified on PDAC-specific NET signature. **(K)** Kaplan-Meier survival analysis of patients with metastatic PDAC stratified as NET-signature-high (groups 1, 2, and 3 in Panel D) and NET-signature-low (group 4 in Panel D). For J and K, Log rank test *P* values, hazard ratios (HR) and 95% confidence intervals (CI) are shown.

While the formation of neutrophil extracellular traps (NETs) by neutrophils has been linked to tumour progression and metastasis^14,15^, here we were interested in determining whether a NET-based gene expression signature could be compiled for identifying PDAC patient outcomes. To stratify PDAC tumours based on relative expression levels of NET-related genes, we assembled a comprehensive RNA-seq dataset encompassing both resectable and metastatic PDAC patient tumours. We reasoned that the use of samples from the primary tumours and from metastases would afford a unique opportunity to define a NET-related gene signature that incorporates clinical samples across the disease continuum. The resectable datasets included PDAC samples from The Cancer Genome Atlas (TCGA) and the Clinical Proteomic Tumour Analysis Consortium-3 (CPTAC-3) databases. RNA-seq data sets for patients with metastatic PDAC were obtained from the Prospectively Defining Metastatic Pancreatic Ductal Adenocarcinoma Subtypes by Comprehensive Genomic Analysis (PanGen, NCT02869802) and the BC Cancer Personalized OncoGenomics (POG, NCT02155621) clinical trials. In total, 369 patient samples were used for initial analysis (TCGA n = 130; CTPAC n = 133; PanGen, n = 81; POG, n = 25).

We first carried out gene-wise co-expression based cluster analysis on this patient cohort using a NET-related gene set which we curated from previous studies in the context of cancer, including TNBC and PDAC^25,26^. Specifically, the NET-related genes were compiled from a previously reported neutrophil gene set^27^, NETosis-related gene sets^28,29^ and eight hub genes identified recently as a NET and EMT-based prognostic model in the context of PDAC^25^.

The optimal clustering solution identified six distinct clusters of PDAC-specific, co-expressed NET-related genes (Fig. 1B). Because of the previously described mechanistic link between NETs and tumour cell-associated CCDC25 and ILK^20^, we investigated whether any of the NET gene clusters correlated with CCDC25 and ILK expression. We found that two of the gene clusters, cluster 5 and cluster 6, were significantly correlated with high levels of *ILK* expression (p = 0. 017), but, interestingly, not with *CCDC25* (Fig. 1B and C). Further analysis of these gene sets using additional established markers of neutrophils, *CD177*^30^ and *OLFM4*^31,32^, further separated the two ILK-associated gene clusters into *OLFM4*-high and *OLFM4*-low groups (Fig. 1B and Extended Data Fig. 1A). Thus, this analysis identified a 31 gene PDAC-specific NET gene expression signature comprising genes from two co-expressed gene clusters that is highly correlated with expression of ILK. The genes present within the NETs signature are as follows: Gene Cluster 5 (n=8 genes: *LDLR, MAPK3, CARD11, GSDMD, SRC, RIPK3, KLF2* and *TICAM1*) and Gene Cluster 6 (n=23 genes: A*CTB, PKM, ACTG1, ACTN4, ENO1, MYH9, CD44, ITGB1, SPP1, TIMP1, ACTN1, AKT2, TKT, F3, AKT1, MFN2, MTOR, MYD88, DNAJB1, CEBPB, DCBLD2, KRT10* and *MAPK7*).

Next, we used this bespoke NET signature gene set to perform sample-wise cluster analysis on the PDAC patient cohort defined above. Consensus clustering of patients using this gene expression signature identified four distinct subgroups of patients (Fig. 1D). While groups 1, 2 and 3 had high levels of expression of the NETs signature genes, group 4 was identified as having low levels of expression of these genes (Fig . 1D). Interestingly, groups 1 and 2 express high levels of NET signature genes, but clearly represent 2 distinct populations (Fig. 1D). Specifically, group 1 patients express high levels of cluster 6 genes and low levels of cluster 5 genes, whereas group 2 patients express high levels of cluster 5 genes (Fig 1D) and low levels of cluster 6 genes. Furthermore, patients in groups 1 and 3 demonstrated significantly higher levels of expression of ILK (p = 2. 2e-11), compared to patients in groups 2 and 4 (Fig. 1E). Levels of expression of CCDC25 were not different amongst the 4 groups (Fig. 1E).

Closer inspection of gene cluster 6 revealed the presence of ITGB1 (Fig. 1D), which is a well-established interactor of ILK^33–35^. Interestingly, this cluster of genes is also highly enriched in genes involved in integrin-actin cytoskeleton organization, including *ACTB* (beta-actin), *ACTG1* (G-Actin), *ACTN1* and *ACTN4* (alpha-actinin)^36^ (Fig. 1D). In addition, other genes established to be involved in ILK signaling, such as *AKT*, *mTOR* and *CD44* are also present in this cluster^34,37^. Significantly, when resectable and metastatic samples were analyzed independently, a higher percentage of patients in group 1 showed overexpression of these ILK-associated genes, compared to group 2 (Fig 1F). Further analysis showed that the expression of many of these ILK associated genes are elevated in groups 1 and 3, compared to group 2 (Fig. 1G and Extended Data Fig. 1B), in addition to elevated protein expression of ITGB1 and CD44, which were quantified by mass spectrometric based proteomics^38^ in a subset of patient samples. (Fig. 1H and Extended Data Fig. 1C).

Gene cluster 5, on the other hand, represents genes involved in cell death, especially necroptosis (*GSMD, RIPK3,* and *MAPK3*) and autophagy (*KLF2*) (Fig. 1D). Here, when resectable and metastatic samples were analyzed independently, a higher percentage of patients in group 2 showed significantly higher levels of expression of these genes, compared to group 1 (Fig. 1I).

To determine whether these patient groups are correlated with differential overall survival, we carried out Kaplan Meier analysis on patient groups 1-3. The results showed that patient group 1, which has a high NET signature and high expression of ILK signaling genes, showed significantly worse overall survival for both resected (p = 0. 023) and metastatic (p = 0. 0064) patient populations, compared to groups 2 and 3 (Fig. 1J). In contrast, patient group 2, which did not represent genes for ILK signaling, but instead consists of genes involved in cell death, had overall better survival outcomes than group 1 and 3 (Fig. 1J).

Finally, analysis of all metastatic PDAC samples showed that NET gene expression signature high patients (groups 1, 2 and 3) survived for a significantly shorter time, compared to NET gene expression signature low patients (group 4) (Fig. 1K).

### Neutrophils and NETs are present in ITGB1, ILK and CCDC25 expressing human PDAC tumours and preclinical xenograft models of PDAC

Our bioinformatics data show that patients with high expression of NET signature genes associated with ILK signaling have poor outcomes and suggests a potential role for NET-mediated ITGB1-ILK signaling in PDAC. To validate that human PDAC tumours harbor neutrophils and NETs, and to determine whether ITGB1 and ILK, as well as CCDC25 and ILK, co-localize in these tissues we analyzed, using multi-color immunofluorescence staining, primary tumour tissue sections from 20 PDAC patients for myeloperoxidase (MPO) and citullinated histone 3 (cit-H3), which are markers neutrophils and NETs, as well as for ITGB1, ILK and CCDC25. We observed regions of co-localization of MPO and cit-H3 in multiple representative tumour samples, demonstrating the presence of NETs in the primary tumours of patients with PDAC (Fig. 2A). We further interrogated these tumours for ITGB1 and ILK, as well as for CCDC25 and ILK. We observed that ILK is highly expressed at the cell surface of PDAC epithelial cancer cells and that ILK and ITGB1 co-localize in these regions (Fig. 2B). While our bioinformatics analysis did not show a correlation with expression of CCDC25, we did observe co-localization of ILK and CCDC25 in the human PDAC tumour tissues (Fig. 2C).

**Figure 2.**
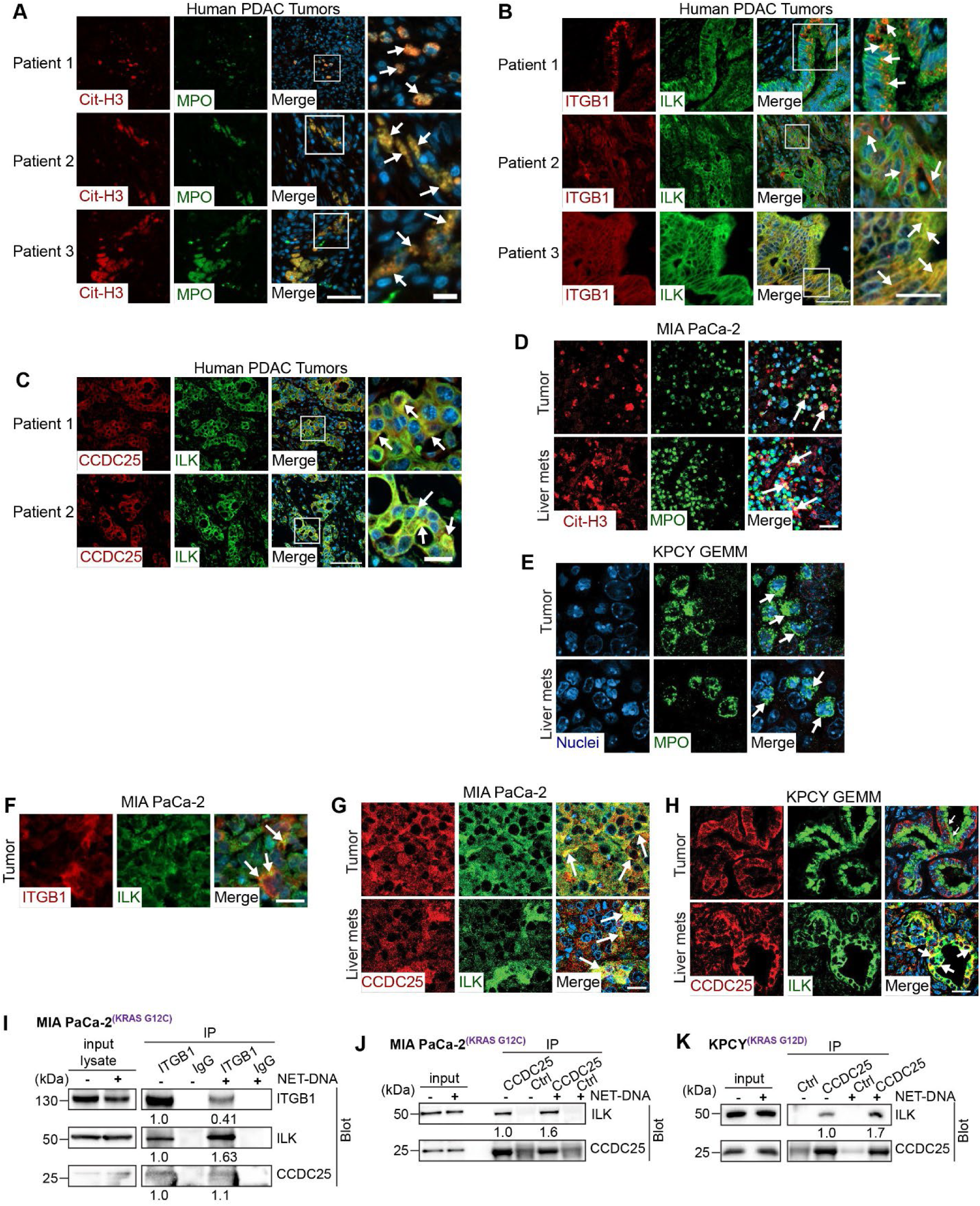
Neutrophils and NETs are present in ITGB1, ILK and CCDC25 expressing human PDAC tumours and preclinical xenograft models of PDAC. **(A)** Human PDAC tumour tissue sections stained by immunofluorescence (IF) for MPO and Cit-H3 to identify neutrophils and NETs (arrows). Scale bar, 50 μm; right panel, 10 μm. Boxes, ROIs shown at higher magnification to the right. Scale bar, 10 μm. **(B, C)** IF staining showing co-localization of (B) ITGB1 and ILK and (C) CCDC25 and ILK (arrows) in representative human PDAC tumour tissue sections. Scale bar, panel B: 50 μm, panel C, 100 μm. Boxes, ROIs shown at higher magnification to the right. Scale bar, 20 μm. **(D)** IF staining of MPO and CitH3 identifying NETs (arrows) in representative tissue sections from primary tumours and liver metastases of MIA PaCa-2 PDAC xenografts. Scale bar, 20 μm. **(E)** IF staining of MPO identifying neutrophils in representative tissue sections from primary tumours and liver metastases of KPCY genetically engineered mouse model of PDAC. **(F)** IF staining showing co-localization of ITGB1 and ILK (arrows) in MIA PaCa-2 xenografts. Scale bar, 20 μm. **(G)** IF staining showing co-localization of CCDC25 and ILK (arrows) in representative primary tumours and liver metastases of MIA PaCa-2 PDAC xenografts. Scale bar, 20 μm. **(H)** IF staining showing co-localization of CCDC25 and ILK (arrows) in representative primary tumours and liver metastases from the KPCY GEMM model. Scale bar, 20 μm. **(I)** Immunoblots showing co-IP of ITGB1, ILK and CCDC25 from MIA PaCa-2 cells cultured with NETs for 24 hours. **(J, K)** Immunoblots showing co-IP of CCDC25 and ILK from (J) MIA PaCa-2 human PDAC cells and (K) KPCY mouse cultured with NETs for 24 hours. Quantification of band intensities is reported below the blots (I, J, K).

To validate these findings further, we interrogated multiple pre-clinical models of PDAC for NETs, ITGB1, ILK and CCDC25. We observed the presence of infiltrating neutrophils and NETs in the both primary tumours and metastases from the MIA PaCa-2 human xenograft (Fig. 2D) and KPCY genetically engineered mouse model (GEMM) (Fig. 2E) of PDAC. Furthermore, ILK is highly expressed in the cancer cells of these models and colocalizes with both ITGB1 and CCDC25 (Fig. 2F-H). Collectively, these data demonstrate, in clinical and experimental PDAC tumours, the presence of the infiltrating neutrophils and NETs, together with the presence of tumour cells expressing ILK in spatial context with ITGB1 and CCDC25.

To determine whether ITGB1, ILK and CCDC25 interact in the context of NET stimulation, we cultured human MIA PaCa-2 cells and mouse KPCY PDAC cells with NETs isolated from DMSO-differentiated, PMA activated HL-60 cells and carried out co-immunoprecipitation (co-IP) assays. We observed that immunoprecipitation of ITGB1 from PDAC cells pulled down both ILK and CCDC25 (Fig. 2I), suggesting that these proteins contribute to a multi-protein complex in PDAC cells. Interestingly, exposure of the cells to NETs resulted in an overall decrease in the levels of ITGB1, but increased amounts of co-immunoprecipitation of ILK, suggesting that the ITGB1-ILK interaction increases with NET stimulation (Fig. 2I). Similarly, immunoprecipitation of CCDC25 from PDAC cells demonstrated that a greater amount of ILK is associated with CCDC25 in cells exposed to NETs (Fig. 2J and K). These data demonstrate, for the first time, a tripartite interaction of ILK with the cell surface receptors, ITGB1 and CCDC25.

### NETs induce PDAC cell matrigel invasion in a NET-DNA, ITGB1 and ILK-dependent manner

Since prior work in the context of breast cancer has demonstrated that NETs stimulate tumour cell migration and invasion^12,15,19,20,39^, we wanted to determine whether NETs induce invasion of PDAC cells. We cultured human MIA PaCa-2 cells and mouse KPCY PDAC cells with NETs and determined the effect of migration and invasion through Matrigel using an incucyte-based kinetic scratch wound assay. We used DMSO-differentiated, PMA activated HL-60 cells as a source of NETs as this system is well-established at providing a standardized source of NETs^40^.

Exposure of PDAC cells to 5-10 µg/mL of NETs, based on NET-DNA concentration, resulted in a significant increase in invasion through Matrigel of MIA PaCa-2 and KPCY cells (Fig. 3A, B and Extended Data Fig. 2A). Exposure of NETs to DNase I, which degrades NET-DNA and thereby inhibits the NET-DNA-CCDC25 interaction^20^, resulted in significant inhibition of NET-stimulated invasion (Fig. 3A, B and Extended Data Fig. 2A). To determine whether the effect of DNase I on invasion by PDAC cells is related to the interaction of NET DNA with CCDC25, we stably transduced MIA PaCa-2 and KPCY cells with doxycycline-inducible shRNA targeting *CCDC25*. Similar to the effect of DNase I, dox-induced shRNA-mediated suppression of *CCDC25* expression reduced NET-induced invasion, compared to control cells (Fig. 3C and Extended Data Fig. 2B), confirming a role of NET DNA-CCDC25 interaction in tumour cell invasion^20^.

**Figure 3.**
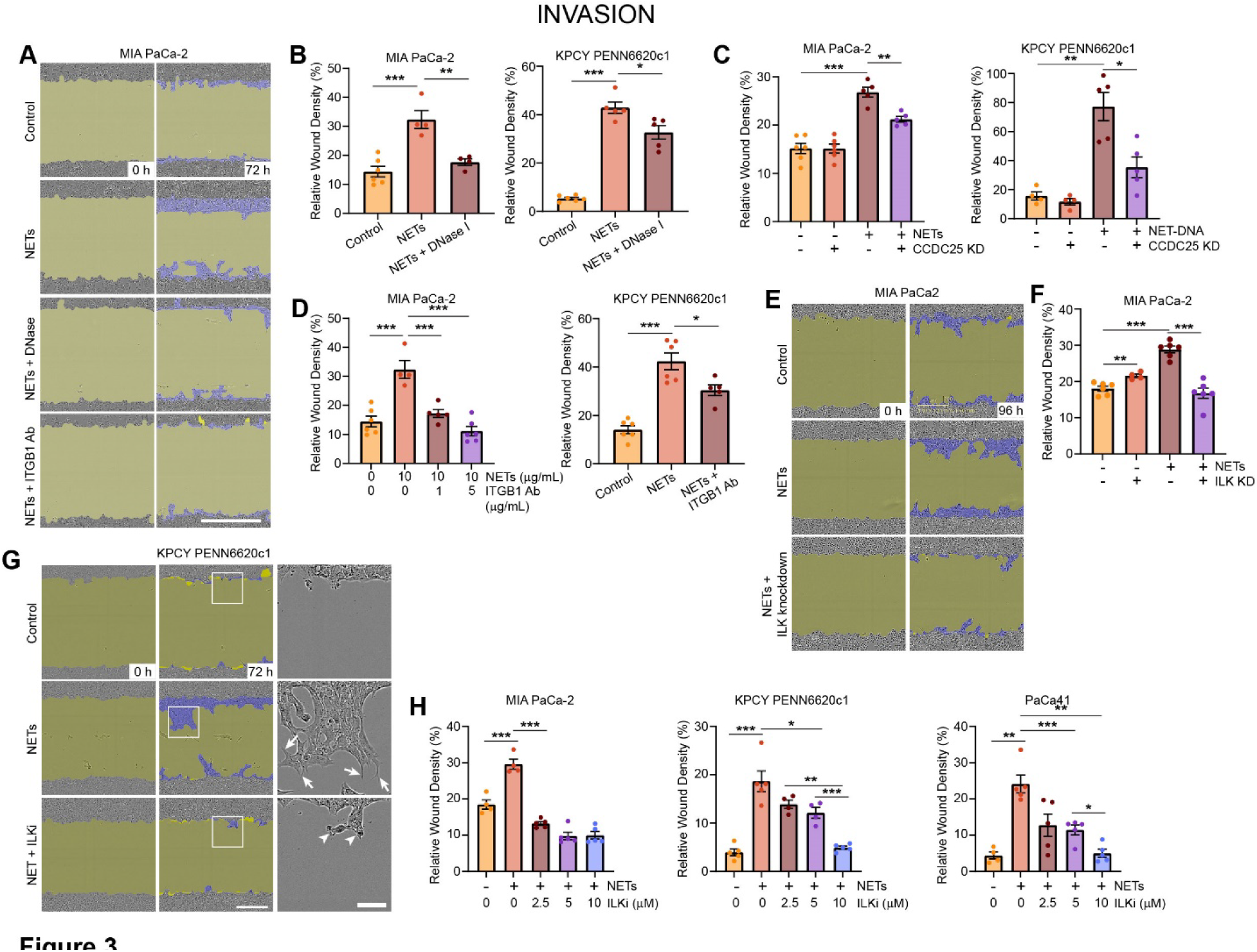
NETs induce PDAC cell matrigel invasion in a NET-DNA, ITGB1 and ILK-dependent manner. **(A)** Time-lapse imaging of invasion through Matrigel by MIA PaCa-2 cells cultured in the presence of 10 µg/mL NETs and exposed to DNase or function-blocking antibodies targeting ITGB1. yellow, wound area; blue, area covered by invading cells. **(B, C, D)** Quantification of NET-induced invasion through Matrigel by human and mouse PDAC cells (B) treated with DNase, (C) following knockdown of CCDC25 expression using dox-inducible shRNA and (D) exposed to function-blocking antibodies targeting ITGB1 for 72 hours. **(E)** Time-lapse imaging showing invasion through Matrigel by MIA PaCa-2 cells following depletion of ILK expression using siRNA and culture in the presence of 10 µg/mL NETs. **(F)** Quantification of NET-induced invasion by cells described in panel E. **(G)** Time lapse images of cells undergoing NET-DNA-induced invasion in the presence of a specific inhibitor of ILK, QLT-0267. Invading cells form robust membrane protrusions (arrows), which are when ILK activity is inhibited (arrowheads). **(H)** Quantification of NET-induced invasion by the indicated cell lines in the presence of the ILK inhibitor. Bars show mean ± s. d. *p < 0. 05, **p < 0. 01, ***p <0. 001, ANOVA (B,C,D, F and H).

However, since the analysis of the clinical data described above (Fig. 1) demonstrated the presence of a group of patients with particularly poor prognosis containing high expression of *ILK* and *ITGB1*, and ILK is known to interact with ITGB1 to control cell, adhesion, migration and invasion^35,36^, we determined the effect of blocking ITGB1 function on NET-induced invasion by PDAC cells. NET-induced invasion of PDAC cells was significantly reduced in the presence of with antibodies targeting the function of ITGB1 (Fig. 3A and D and Extended Data Fig. 2C). Blocking ITGB1 function also inhibited NET-induced migration by MIA PaCa-2 cells (Extended Data Fig. 3A).

Since both CCDC25 and ITGB1 are known to interact with ILK, we next determine the effect of inhibiting ILK on NET-induced invasion using both genetic and pharmacologic strategies. siRNA-mediated depletion of *ILK* expression resulted in significant inhibition of NET-stimulated migration (Extended Data Fig. 3B) and invasion (Fig 3E, F and Extended Data Fig. 2D) of MIA PaCa-2 cells. Furthermore, pharmacologic inhibition of ILK activity, using an established ILK inhibitor, QLT0267^37,41,42^ resulted in abrogation of NET-induced invasion in a dose dependent manner across multiple PDAC cell lines, including in PDX-derived PaCa41 cells (Fig. 3G, H and Extended Data Fig. 2E).

Pharmacologic inhibition of ILK had a modest effect on basal levels of invasion by the MIA PaCa-2 cells and had no effect on basal invasion levels in KPCY or PaCa41 cells (Extended Data Fig. 2F). The ILK inhibitor also reduced basal (Extended Data Fig. 3C) and NET-induced migration (Extended Data Fig. 3D) by MIA PaCa-2 cells in a dose-dependent manner. Collectively, these data demonstrate that NETs induce migration and invasion by PDAC cells and that this increased motility is mediated by an ITGB1/CCDC25/ILK signaling axis.

### Epithelial to Mesenchymal Transition (EMT) is a hallmark of NETosis in PDAC tumours

Cellular plasticity is now considered a requisite characteristic of cancer cells capable of metastasis. EMT is a key process that underlies cancer cell invasive and migratory abilities and is critical for cancer progression and metastasis.

To identify differential enrichment of cellular pathways between the prognostic NET signature groupings identified in our bioinformatics analysis (Fig. 1), we next performed comprehensive gene set enrichment analysis (GSEA) on genes differentially expression between patient groups 1 and 2 (Fig. 4A). This analysis revealed EMT to be the most highly enriched pathway among genes up-regulated in the poor outcomes patient group 1 relative to the better outcomes group 2 (Fig. 4A). Gene sets related to hypoxia and inflammatory response were also highly enriched among the upregulated genes in the group 1 patients (Fig. 4A), both of which are implicated in aggressive PDAC tumour cell behavior. Other gene sets significantly enriched among genes up-regulated in group 1 patient tumors included neutrophil degranulation and integrin cell surface interactions (Fig. 4A), which are pathways that would be expected given the NET-mediated regulation of the ITGB1-ILK interaction that we have uncovered.

**Figure 4.**
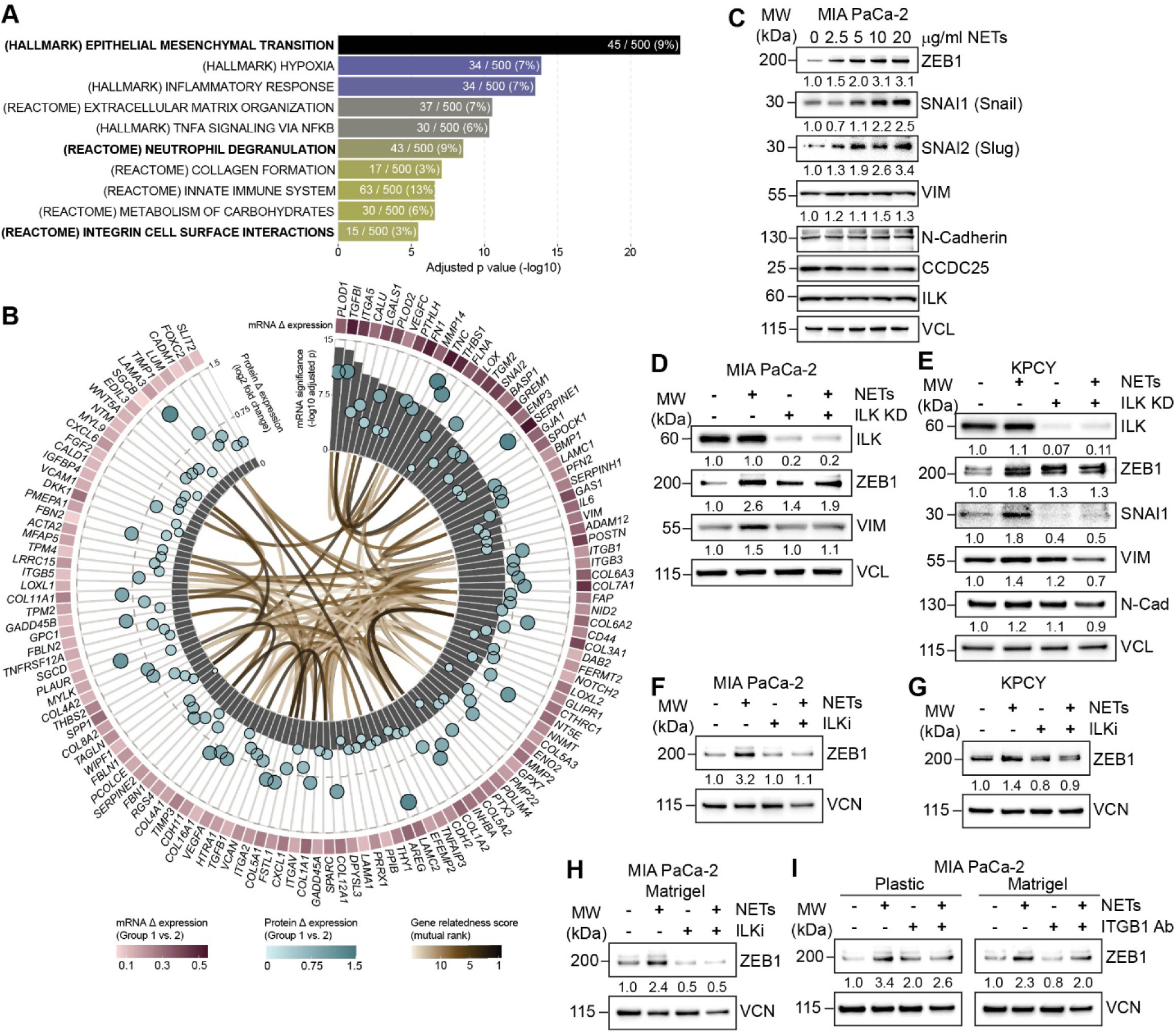
Epithelial to Mesenchymal Transition (EMT) is a hallmark of NETosis in PDAC tumours. **(A)** GESA of differentially expressed genes in group 1 (poor outcome) versus group 2 (better outcome). Bar plot showing pathways upregulated in patient group 1 (poor prognosis) versus patient group 2 (better prognosis). **(B)** Analysis of Hallmark EMT genes significantly upregulated in patient group 1 compared to patient group 2. Grey bars, strength of significance of gene dysregulation; Red squares, magnitude of differential expression; Teal bubbles, difference in protein levels for each gene; Brown ribbons, gene-gene relatedness based on GeneFriends analysis. **(C)** Western blot analysis of the indicated EMT markers in MIA PaCa-2 cells cultured in the presence of increasing concentrations of NETs for 24h. **(D, E)** Western blot analysis of NET-induced EMT markers in (D) MIA PaCa-2 and (E) KPCY cells in response to knockdown of ILK expression. **(F, G)** Western blot analysis of NET-induced ZEB1 expression in (F) MIA PaCa-2 and (G) KPCY cells in response to pharmacologic inhibition of ILK. **(H)** Western blot analysis of NET-induced ZEB1 expression in MIA PaCa-2 cells cultured on Matrigel and exposed to the ILK inhibitor. **(I)** Western blot analysis of NET-induced ZEB1 expression in MIA PaCa-2 cultured the indicated substrates in the presence of function-blocking Abs against ITGB1. Quantification of band intensities is reported below the blots (C-I).

Next, we further interrogated genes belonging to the Hallmark EMT gene set that were significantly upregulated in patient group 1 (poor prognosis) compared to patient group 2 (better prognosis) (Fig. 4B). We found significant upregulation of expression of several EMT genes in patient group 1, many of which also showed overexpression at the level of protein (Fig. 4B). Furthermore, the expression of many of these genes are highly related, based on the GeneFriends^43^ mutual rank score, suggesting strong co-expression of these genes (Fig. 4B). Collectively, these data identify EMT has a critical pathway upregulated in PDAC patients that are NET signature high, ILK signaling high and subject to poor outcome.

Since our bioinformatics analysis suggested that EMT is a critical process in patients identified as NET signature high and ILK signaling pathway high, and who have poor outcomes, we next wanted to determine whether NETs stimulate EMT in PDAC cells. First, we incubated PDAC cells with increasing concentrations of NETs and analyzed the protein levels of transcriptional regulators of EMT. We observed, by western blot analysis, that exposure to NETs results in robust, dose-dependent increases in EMT markers SNAI1, SNAI2 and, in particular, ZEB1 (Fig. 4C and Extended Data Fig. 3E). We further observed a dose-dependent increase in the mesenchymal marker, vimentin. Next, since previous studies have shown that ILK promotes EMT in the context of cancer^44–46^, we knocked down expression of ILK using dox-inducible shRNA and assessed the impact on these EMT markers. In both the human and mouse PDAC cells, exposure to NETs increases the levels of ZEB1 and vimentin, and silencing ILK abrogates this NET-induced increase (Fig. 4D and 4E). Silencing of *ILK* expression further inhibited expression of Snail and N-cadherin in the KPCY cells (Fig. 4E).

Furthermore, pharmacologic inhibition of ILK activity also abrogated the NET-induced increase in expression of ZEB1 in both human and mouse cells (Fig. 4F and 4G), and the effect was similar when cells were grown on plastic (Fig. 4F, G) or Matrigel (Fig. 4H), confirming that inhibition of ILK reduces EMT markers in these cells.

Finally, we found that culturing PDAC cells on either plastic or Matrigel in the presence of function-blocking antibodies targeting ITGB1 phenocopied the inhibition of ZEB1 expression observed with inhibition of ILK (Fig. 4I), further indicating that NETs induce EMT in PDAC cells through an ITGB1-ILK signaling axis.

### NETs induce pseudopodia-like protrusions in an ILK dependent manner and signal through GSK3β and Rac1/Cdc42 to induce invasion

Our data clearly show that NETs induce invasion by PDAC cells and that ILK is an important contributor to NET-stimulated invasion. To further investigate the mechanism of ILK-mediated NET-induced invasion, we investigated the colocalization of ILK with paxillin, a marker of focal adhesions, and cofilin, a marker of pseudopodia-like protrusions involved in metastatic colonization^47^. Given our results demonstrating increased invasion through matrigel and the potential role of ITGB1 in this process, we performed these studies using PDAC cells grown on Matrigel and fibronectin.

Exposure of PDAC cells to NETs induced dramatic changes in cellular phenotype. In the presence of NETs, cells cultured on Matrigel (Fig. 5A, C) and fibronectin (Fig. 5B, D) increased in overall size and extended pseudopodia-like protrusions (PLPs), suggesting the acquisition of motility. Paxillin co-localized with ILK at focal adhesions in cells plated on Matrigel (Fig. 5A) fibronectin (Fig. 5B). Exposure of cells to NETs resulted in redistribution of these proteins such that they showed dramatic co-localization with PLPs in NET-induced cells.

**Figure 5.**
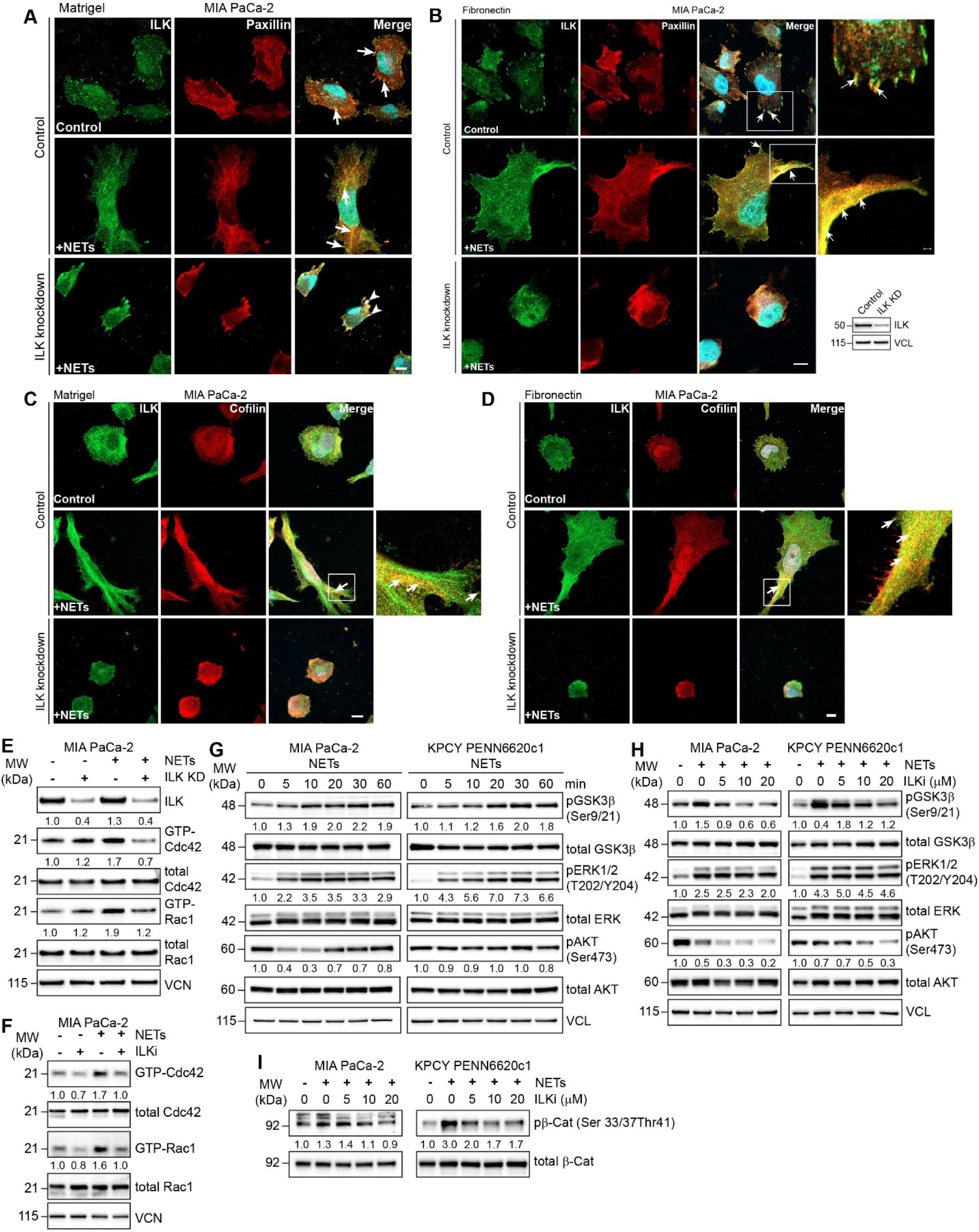
NETs induce pseudopodia-like protrusions in an ILK dependent manner and signal through GSK3β and Rac1/Cdc42 to induce invasion. **(A, B)** IF staining showing co-localization (arrows, yellow) of ILK and paxillin in control and ILK-depleted MIA PaCa-2 cells cultured on (A) Matrigel and (B) fibronectin with 20 μg/mL NETs for 24 hours. Boxes, ROIs shown at higher magnification in right panels. Scale bar = 10 µm. **(C, D)** IF staining showing co-localization (arrows, yellow) of ILK and cofilin in control and ILK-depleted MIA PaCa-2 cells cultured on (C) Matrigel and (D) fibronectin with 20 μg/mL NETs for 24 hours. Boxes, ROIs shown at higher magnification in right panels. Scale bar = 10 µm. **(E)** Immunoblot analysis showing GTP-bound and total Rac1 and Cdc42 levels in control and ILK-depleted MIA PaCa-2 cells cultured with or without 10 μg/mL NETs for 24 hours. **(F)** Immunoblot analysis showing GTP-bound and total Rac1 and Cdc42 levels in MIA PaCa-2 cells cultured with or without 10 μg/mL NETs and the ILK inhibitor. **(G)** Western blots showing levels of phosphorylation of the indicated proteins in NET-stimulated PDAC cells at the indicated time points. **(H)** Western blots showing levels of phosphorylation of the indicated proteins in NET-stimulated PDAC cells response to increasing concentrations of the ILK inhibitor. **(I)** Levels of phosphorylation of β-catenin in PDAC cells cultured as described in panel H. Quantification of band intensities is reported below the blots (E-I).

Dox-inducible knockdown of *ILK* expression inhibited the ability of PDAC cells to develop membrane protrusions in the presence of NETs on either substrate (Fig. 5A-D), demonstrating a requirement of ILK expression and function.

We also assessed the localization of cofilin, a marker of PLPs^47^, and ILK in the context of NET stimulation. In control cells plated on either Matrigel or fibronectin, ILK was distributed both within the cytoplasm and at the plasma membrane where it was situated within focal adhesions, while cofilin was diffusely distributed throughout the cells (Fig. 5C, D). Upon exposure to NETs, the cells developed protrusions at which ILK and cofilin were enriched and colocalized (Fig. 5 C, D). Cells in which dox-inducible shRNA targeting ILK was used to suppress ILK expression remained round and did not develop protrusions in response to exposure to NETs (Fig. 5C, D).

Since NETs stimulate both membrane protrusions and invasion in PDAC cells, we wanted to understand the potential mechanism. Since the GTPases, Rac and CDC42 have been shown to regulate cellular migration and invasion^48,49^, we investigated whether NETs can stimulate the activities of these GTPases in PDAC cells. Exposure to NETs increased GTP loading of both Rac1 and Cdc42, indicating an increase in Rac1 and Cdc42 activity (Fig 5E). Inhibition of ILK activity with QLT-0267 reduced the levels of both GTP-Rac1 and GTP-Cdc42, indicating inhibition of this signaling axis (Fig. 5F).

Since our data demonstrate that NETs induce EMT in PDAC cells and stimulate activation of Rac and Cdc42 to induce invasion, we wanted to further interrogate additional signaling pathways know to be associated with ILK-mediated EMT, such as GSK3β and β-catenin^34,45,46^. Exposure of human and mouse PDAC cells to NETs results in a time-dependent increase in the phosphorylation of GSK3β as well as ERK (Fig. 5G). Pharmacologic inhibition of ILK activity using increasing concentrations of the ILK inhibitor, QLT-0267, results in dose-dependent inhibition of GSK3β in both cell lines (Fig. 5H). Phosphorylation of Akt on Ser473 was also abrogated by the ILK inhibitor in a dose-dependent manner, confirming that the inhibition of NET-induced GSK3β is ILK dependent (Fig. 5H).

Similarly, phosphorylation of β-catenin increased in response to the exposure of both human and mouse PDAC cells to NETs, and this increase in phosphorylation was dose-dependently inhibited in response to incubation with the ILK inhibitor (Fig. 5I).

We also examined the effect of NET stimulation on cell growth, since previous studies in breast cancer cells^20^ have shown that NETs not only stimulate migration and invasion, but also cell growth which may further contribute to metastatic colonization. Exposure of PDAC cells in 3D MiaPaCa2 spheroids, MIA PaCa2 showed increased size of spheroids in the presence of NETs, (Extended Data Fig. 4A). NET-induced growth of spheroids is significantly reduced in the presence of DNase I (Extended Data Fig. 4A, B), the ILK inhibitor (Extended Data Fig. 4A, B) or ITGB1 antibodies (Extended Data Fig. 4C). Thus, in addition to impacting migration and invasion, NETs induce growth of PDAC cells in 3D and this effect can be inhibited by interfering with CCDC25, ITGB1 and ILK activity.

### Inhibition of ILK suppresses NET-induced metastasis

Our data demonstrate that the ITGB1-ILK signaling axis is a critical contributor to NET-induced PDAC tumour cell EMT and invasion, suggesting that targeting this signaling in vivo may provide a novel therapeutic avenue for reducing metastasis in inflammation driven PDAC.

To test this, we used an experimental metastasis model wherein we injected MIA PaCa-2 PDAC cells expressing both luciferase and dox-inducible shRNA targeting *ILK* into the tail vein of NSG mice that were administered LPS intranasally immediately prior to tumour cell injection, as well as at days 3 and 6 post-tumour cell injection to induce lung inflammation (Fig 6A). Mice were administered dox in the drinking water (or drinking water without dox as a control) following cell injection (Fig 6A) to induce shRNA-mediated knockdown of ILK protein expression (Fig. 1B) and metastasis was monitored using bioluminescence imaging. We observed that knockdown of *ILK* expression in the tumour cells in the context of LPS-stimulated lung inflammation dramatically reduced overall metastatic burden at 42 days post cell injection, compared to control animals (Fig. 6C). Quantification of the bioluminescence signal demonstrated a significant reduction in metastatic burden, as measured by total flux, with ILK knockdown, compared to controls (Fig. 6D).

**Figure 6.**
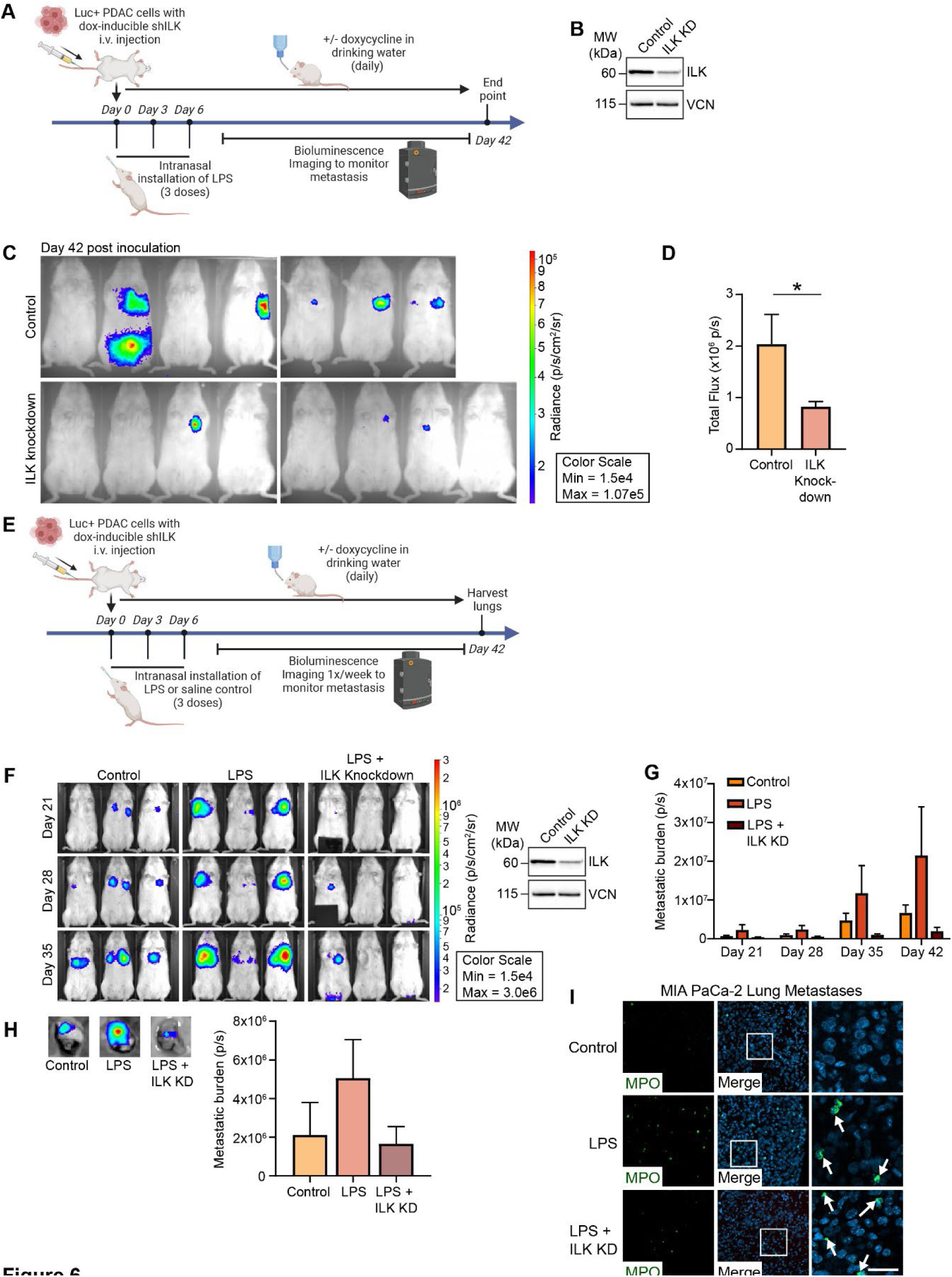
Inhibition of ILK suppresses NET-induced metastasis. **(A)** Schematic showing the in vivo experimental design. **(B)** Western blot showing dox-inducible knockdown of ILK expression in MIA PaCa-2 Luc+ dox-inducible shILK cells. **(C)** Bioluminescence images showing metastatic burden in control mice and mice treated with dox to induce knockdown of ILK expression. **(D)** Quantification of bioluminescence signal in panel C (n = 7 to 8/group). Bars show mean ± s. e. m. *p < 0. 05, *t-*test. **(E)** Schematic showing the in vivo experimental design. **(F)** Bioluminescence images showing lung metastatic burden in control mice and mice administered LPS intranasally, with or without dox treatment at the indicated time points. **(G)** Quantification of bioluminescence signal in the lungs at the indicated time points (n = 5 to 6/group). **(H)** Bioluminescence images of representative lung metastases from the indicated treatment groups at day 42 post cell injection and quantification of metastatic burden. (n = 5 to 6/group). **(I)** IF staining of MPO in lung metastases from mice treated as indicated in panel F at 42 days after injection of MIA PaCa-2 tumour cells. Scale bar, 100 μm, 25 μm (right panel).

We then further determined whether LPS-driven lung inflammation was indeed driving increased metastasis and, if so, whether the increased metastasis was being inhibited by knockdown of ILK expression. For these studies, mice were administered LPS, or saline as a control, intranasally immediately prior to injection of the tumour cells and at days 3 and 6 post tumour cell injection (Fig. 6E). MIA PaCa-2 Luc+ dox-inducible *ILK* shRNA tumour cells were inoculated through the tail vein and lung metastatic burden was monitored by bioluminescence imaging. We found a clear, time-dependent increase in metastatic burden in the lungs of mice administered LPS, compared to control mice administered saline (Fig. 6F, G), demonstrating that LPS-induced inflammation and neutrophil recruitment results in increased metastasis in this model. Knockdown of ILK expression in the PDAC tumour cells results in a dramatic reduction in lung metastatic burden at all time points examined (Fig. 6F, G). Ex vivo imaging of metastatic burden in the lungs at day 42 post cell injection also demonstrated increased metastases in LPS-treated animals, and reduced metastatic burden in the lungs of mice in which ILK was depleted from tumour cells (Fig. 6H). Immunofluorescence for MPO in lung tissue sections from the mice at end-stage showed that mice treated with LPS had increased numbers of neutrophils localized within the metastatic foci, compared to control mice that showed neutrophils largely at the periphery of the metastases (Fig. 6I). While the lung metastatic burden in the mice treated with LPS and injected with *ILK* KD cells was reduced (Fig. 6F-H), we did observe that neutrophils were still present, albeit somewhat less numerous, within the metastases of this group (Fig. 6I), in keeping with the LPS-mediated stimulation of inflammation in this group.

## Discussion

It is now well established that the tumour microenvironment influences various aspects of cancer progression to metastasis, the root cause of cancer mortality. The tumour microenvironment can, in turn, be influenced by the host state, which can govern the immune cell repertoire of the microenvironment. Poor patient outcomes are also influenced by the immune suppressive microenvironment, which in turn is regulated through hypoxia, vascularization, extracellular matrix, establishment of a pre-metastatic niche^50^, and inflammation, The latter is known to influence tumour behaviour, and the inflammatory cells such as monocytes, macrophages and neutrophils play significant roles in immune-suppression. A recent detailed study on the spatial architecture of myeloid and T cells demonstrated that the ratios of these cells play a critical role in immune evasion and clinical outcome in lung cancer^51^. In particular tumour associated neutrophil infiltration identified a subset of tumours which had increased metastatic propensity and shorter disease-free survival^51^.

While neutrophil infiltration of tumours and neutrophil to lymphocyte ratios (NLR) are associated with poor patient outcomes^10,11,24^, the molecular basis of how neutrophils influence tumour cell behaviour is less well understood. Recent reports have suggested that inflammatory stimuli-driven NETosis, leading to the formation of neutrophil extracellular traps (NETs) promote aggressive tumour cell phenotypes such as migration/ invasion and metastatic colonization^12,13,15^. NETs have also been implicated in “awakening” dormant tumour cells within the circulation to extravasate and colonize distant organs^14^. NETs are complex structures composed decondensed genomic DNA, histones and proteases^17,18^, and the precise mechanisms of how NETs influence tumour cell phenotype are still under investigation.

A recent study demonstrated a role of the modified DNA component of NETs, and demonstrated in breast cancers that the binding of NET DNA to a cell surface protein, CCDC25 plays a critical role in inducing tumour cell migration/invasion and metastasis formation by recruiting integrin-linked kinase/beta –parvin^20^, which are established effectors of cell-cytoskeleton organization and dynamics^34,47^.

Here, we have investigated the role of NETosis and NETs in pancreatic adenocarcinoma (PDAC) progression to metastasis. PDAC is a very aggressive cancer with a complex, immunosuppressive tumour microenvironment. While previous studies have determined that inflammation-driven neutrophil infiltration and NETosis are found in PDAC tumours, it is unclear whether NETosis and NETs can be used to identify PDAC patient out comes and prognosis, and the downstream PDAC-tumour cell effectors of NET-tumour cell interactions that drive invasion and metastasis have not been identified for potential therapeutic targeting to prevent and inhibit NET-driven metastasis.

Here, by utilizing bioinformatics approaches in PDAC patient tumour samples composed of primary resectable tumours as well as metastases, together with cell biological approaches and in vivo tumour models, we now identify NET gene expression signatures that provide information on patient outcomes. Importantly, the analyses in PDAC cohorts identify NET signature-associated expression of genes that reveal potential molecular pathways important in the NET-driven phenotype. Specifically, we find that NET signatures that are associated with high Integrin-Linked Kinase (*ILK*), beta1 integrin (*ITGB1*), and high actin cytoskeleton dynamic genes identify PDAC patients with poor overall survival. The NET-signature high, ILK high tumours are significantly associated with EMT and have increased expression of genes involved in integrin/ILK mediated cytoskeleton organization and cell survival and growth, all of which contribute to tumour cell aggressive behaviour. Furthermore, gene expression signatures identified here from the PDAC RNA-seq data sets are comprised of genes largely associated with tumour cells and do not contain genes encoding for cytokines, MPO or PADI4, which are associated with tumour microenvironment cells such neutrophils, providing support for a our analysis of the effect of NETs on tumour cells.

We have gone on to validate that PDAC tumours and metastases harbour neutrophils and NETs, by immunostaining patient primary tumours and metastases, as well experimental tumours and metastases from human PDAC cell xenogafts and KPCY mouse transgenic tumours and metastases, with antibodies against MPO (neutrophils) and citrulinated H3 (NETs). The data clearly demonstrate that not only do PDAC tumours and metastases have significant numbers of intra-tumoural neutrophils, but that a substantial number undergo NETosis and produce NETs. Counter staining of adjacent serial sections also demonstrated that these NETs surround tumour cells expressing ITGB1 and ILK.

Based on these data we set out to determine the molecular basis of NET-mediated stimulation of cell migration, invasion, growth and, especially, metastasis. Utilizing PDAC cells in culture, we have shown that ILK forms a complex with both ITGB1 and CCDC25, as determined by immunostaining and co-immunoprecipitation, and that NETs stimulate tumour cell invasion and migration in NET DNA, CCDC25, ITGB1 and ILK-dependent manner. Furthermore, NETs promote a dramatic morphological switch to a highly motile and invasive phenotype that is accompanied with increased formation of ILK and cofilin containing pseudopodia like protrusions (PLPs), as well as activation of the GTPases Rac and cdc42. These structures have previously been demonstrated to require ILK-cofilin, and Rac/cdc42 activation signaling pathways and to play critical roles in breast tumour cell metastatic colonization^47^. Additionally, ITGB1-ILK complex also been implicated in metastatic colonization of breast cancer cells by promoting LICAM-mediated activation of YAP^52^, a downstream effector of ILK^53^.

Mechanistically we show for the first time that NETs stimulate integrin and ILK-dependent expression of regulators of EMT, such as SNAI1, SNAI2 and ZEB1. Furthermore, NETs induce phosphorylation of GSK-3 and Akt in an ILK dependent manner. GSK-3 inactivation by phosphorylation regulates beta-catenin driven EMT, and also stabilizes snail protein levels by preventing its proteasome-mediated degradation^54,55^. Phosphorylation of Akt regulates cell survival and growth through mTOR^56,57^, a NET signature associated gene (Fig. 1).

Finally, we demonstrate that metastatic colonization of human and mouse PDAC cells after tail vein injection is stimulated by LPS-induced inflammation and associated NETosis. Importantly, inhibition of expression of ILK, using doxycyclin inducible ILK shRNA, significantly suppresses lung colonization in both models, demonstrating that inflammation and NET-driven metastasis requires ILK. While we were able to demonstrate the presence of neutrophils within the lung metastases of mice treated with LPS and show that neutrophils are still present in the metastases formed by *ILK* KD cells, these analyses were carried out here on lungs with end-stage disease. It is important to note that the contribution of neutrophils and NETosis may also occur within the circulation and/or the lung vasculature in a temporal manner. The precise location and timing of the neutrophil and NET-tumour cell interactions will require future analyses using technologies such as intravital microscopy.

Collectively, our data identify novel NET associated gene expression signatures that can be used to identify patients susceptible to neutrophil/NET mediated metastasis and poor outcome. While previous studies have shown the presence of NETs in metastases, an outstanding question has been whether neutrophils and NETs accumulate in primary tumours as well. Our bioinformatics analysis and direct immune-localization experiments suggest that in PDAC, NETosis can be a prominent feature in primary tumours. These findings suggest that analysis of primary tumour tissues for NETs and NET gene signatures could potentially provide critical information on patient outcome and metastasis formation.

Significantly, we have identified a NET signature associated, targetable, signaling axis: Integrin-ILK-EMT in PDAC tumours. While our results implicate ILK as a central player in NET-induced PDAC metastasis, our data point to a number of other targets in NET-induced signaling axis identified in this study (Fig 7).

**Figure 7.**
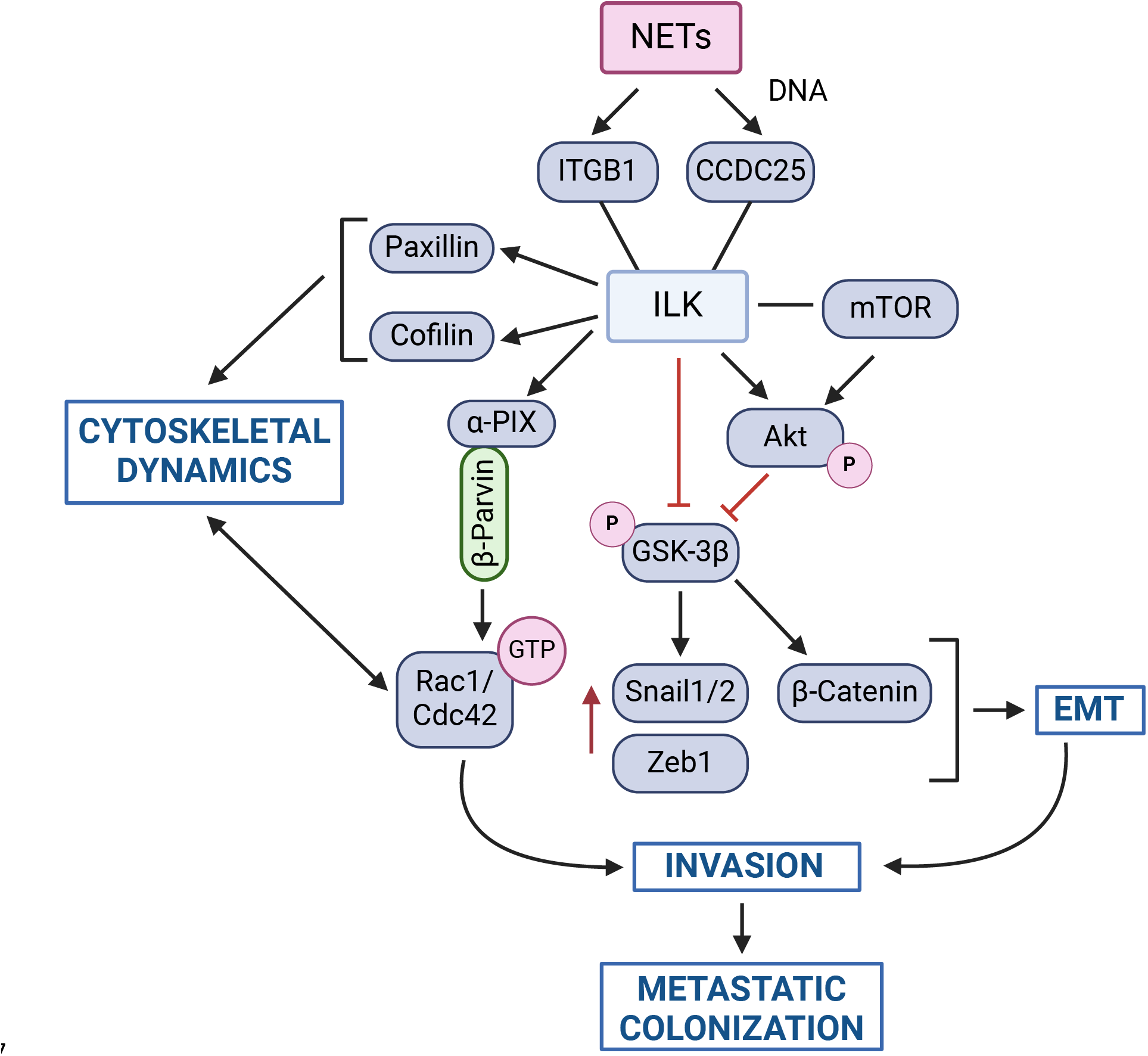
Summary schematic showing the role of NET-induced ILK signaling in PDAC invasion and metastasis.

Finally, targeting the downstream, PDAC tumour cell-associated effectors of NETosis and NETs, identified here, may be more advantageous and specific for the neutrophil/NET promotion of metastasis, than targeting NET formation per se, since neutrophils and NETs have a critical physiological role in fighting infections.

## Methods

### Cells

Human pancreatic cancer cells (MIA PaCa-2) were obtained from Don Yapp and Sylvia Ng (BC Cancer Research Centre, Vancouver, Canada) and cultured as previously described ^58,59^. The cell lines were maintained in Dulbecco’s modified Eagle’s medium (DMEM; Life Technologies) supplemented with 10% fetal bovine serum (FBS; Life Technologies). The congenic PDAC tumour cell clone PENN6620c1 was established in culture from the C57BL/6 KPCY genetically modified mouse model, was provided by Ben Stanger (University of Pennsylvania, Philadelphia, USA) and was cultured as previously described^60^. The PENN6620c1 clone was maintained in DMEM supplemented with 10% FBS. The PaCa41 human patient derived xenograft (PDX) cell line PaCa41 was established from a patient tissue fragment and was provided by Daniel Renouf (BC Cancer). PaCa41 cells were maintained in DME/F-12 medium supplemented with 0. 25 mg/mL bovine serum albumin (BSA) and 5 mg/ml glucose (Fisher Scientific), 1x insulin-transferrin-selenium (ITS) and 25 μg/mL bovine pituitary extract (BPE; Life Technologies), 40 ng/mL epidermal growth factor (EGF; Sigma Aldrich), 5 nM 3,3,5-tri-iodo-L-thyronine and 1 μg/mL dexamethasone (Sigma Aldrich), 100 ng/mL cholera toxin, (Molecular Probes, ThermoFisher), 1. 22 mg/mL nicotinamide (Sigma Aldrich), 5% Nu-serum IV culture supplement (Corning), 100 U/mL penicillin, 100 μg/ml streptomycin and 500 μg/mL amphotericin B (Life Technologies). All cells were grown at 37°C in a humidified atmosphere containing 5% carbon dioxide (CO_2_) and were tested for mycoplasma contamination using the LookOut Mycoplasma PCR detection kit (Sigma; cat no. MP0035). The MIA PaCa-2 cell line was authenticated using short tandem repeat STR DNA profiling by a commercial testing facility (Genetica, Burlington, NC, USA). The KPCY clones have been authenticated as described. ^60^ The PaCa41 cell line was authenticated by the source laboratory. QLT-0267, an inhibitor of ILK activity, was obtained from Quadralogic Technologies Inc (QLT) and used at the concentrations indicated in the text and figures. siRNA for ILK (Qiagen, cat no GS3611 Flexitube Gene Solution) and was used according to manufacturers recommendations.

### Animal studies

All experimental animal procedures were carried out at the BC Cancer Animal Resource Centre (ARC) in accordance with protocol A21-0263 approved by the institutional Animal Care Committee (ACC) at the University of British Columbia, Vancouver. The studies are compliant with all relevant ethical regulations regarding animal research. All mice were housed in ventilated cages in a pathogen-free, environment-controlled room at 19-21°C. The relative humidity ranged between 40% and 70% and a photoperiod of 12 hours light and 12 hours darkness was provided. Food and water were provided ad libitum.

For experimental metastasis studies, MIA PaCa-2 Luc+ dox-inducible shRNA cells (2. 0 x 10^6^ cells/animal, suspended in 100 μL sterile saline) were injected intravenously through the tail vein. Mice were administered lipopolysaccharide (LPS) (0. 25 mg/ml, 50 μL/animal), or equal volumes of saline as a control, by intranasal instillation on days 0, 3, and 6 post injection of tumour cells to induce lung inflammation. On day 0, administration of LPS or saline was performed immediately prior to injection of the tumour cells. To induce the shRNA targeting ILK, mice were administered dox (1 mg/mL) in drinking water containing 1% sucrose ad libitum starting immediately following injection of the tumour cells and continuing for the duration of the study. Control animals were similarly administered drinking water containing 1% sucrose. Metastasis was monitored non-invasively using bioluminescence imaging (BLI). To measure metastatic burden, mice were administered d-luciferin (Promega) at a dose of 150 mg/kg by intraperitoneal injection and imaged using a Perkin Elmer Lumina S3 instrument as described by us previously^59^.

Lung tissues were harvested, fixed in formalin and embedded in paraffin for downstream analyses by immunohistochemistry (IHC). For ex vivo imaging of lung metastatic burden, lungs were harvest from mice administered luciferin as described above and imaged prior to formalin fixation. Body weights were recorded routinely during the course of the study as a measure of animal health. A clinical observation and health monitoring record was also maintained for each individual animal throughout the study. The investigators were not blinded to the identity of the study groups and the number of animals per group was chosen based on data from previous studies^61,62^.

Immunocytochemical analyses of NETs, ITGB1, CCDC25 and ILK in the MIA PaCa-2 xenograft model and the *Kras^G12D^/Pdx1-Cre/p53/Rosa^YFP^* genetically engineered mouse model (GEMM) of PDAC used archived tissues from previously published studies^59,63^.

### Human participants – Resectable PDAC sequencing datasets

RNA-seq data (transcripts per million (TPM); hg19) for resectable PDAC samples from the TCGA database (n=130) were downloaded and log10-transformed, and samples were filtered as described previously^64^. Additional public RNA-seq data for PDAC patient samples from the CPTAC-3 cohort (resectable disease; n=133) were downloaded from the GDC data portal (https://portal.gdc.cancer.gov) on May 27, 2021 (HTSEQ; FPKM) and log10-transformed.

### Human participants – Metastatic tumour datasets

Sequencing data for patients with metastatic PDAC (n=106) was obtained from two prospective studies: Prospectively Defining Metastatic Pancreatic Ductal Adenocarcinoma Subtypes by Comprehensive Genomic Analysis (PanGen, NCT02869802; n=81) and the BC Cancer Personalized OncoGenomics program (POG, NCT02155621; n=25). All PanGen and POG data were sequenced at the Canada’s Michael Smith Genome Sciences Centre in Vancouver, Canada. Patients participating in PanGen and POG studies were enrolled as previously described^38,65^. PanGen and POG studies were approved by the University of British Columbia Research Ethics Committee (REB# H12-00137, H14-00681, H16-00291) and conducted in accordance with international ethical guidelines. Written informed consent was obtained from each patient prior to molecular profiling. All sequencing data were housed using a secure computing environment. The PanGen and POG clinical trials are not directly linked to a specific treatment, but rather aim to assess response to genomics-guided therapy, with treatments selected at the discretion of the treating oncologist. RNA-sequencing (RNA-seq) was performed on metastatic patient tumour samples with a target depth of 200 million reads. RNA-seq reads were trimmed to 75bp and aligned (GRCh37-lite) using STAR v2. 7. 3^66^, with parameters: -chimSegmentMin 20 -outSAMmultNmax 1 - outSAMstrandField intronMotif -outFilterIntronMotifs RemoveNoncanonical. RNA-seq duplicate reads were marked using PicardTools v2. 17. 3. Raw reads counts were assigned to Ensembl 75 genes using Subread v1. 4. 6^67^, normalized for library depth and gene size (RPKM) and log10-transformed.

### RNA-seq batch correction

RNA-seq data was corrected for cohort-specific batches (PanGen/POG, TCGA and CPTAC-3) using an empirical Bayesian approach^68^. Principal component analysis (PCA) of the top 10% most variable genes was used to confirm alleviation of inter-sample batch effects after correction.

### Clustering analysis

Normalized gene expression values were converted to z-scores prior to clustering using all samples combined (n=369). Genes previously implicated in NET pathway expression in other cancers were curated from studies of triple-negative breast cancer (n=136 genes)^26^ and PDAC (n=8 genes)^25^ and used for gene-gene clustering to identify PDAC-specific co-expressed NET signature genes. From this curated gene panel, a total of 123 NET-related genes were present within the RNA-seq datasets and comprised the NET-related gene set used for gene-gene cluster analysis (Extended Data Table 1).

Genes identified from this analysis comprised a PDAC NET-signature gene set (n=31 genes) and were then used as input for subsequent sample-sample clustering. These genes were identified from two co-expressed clusters: Gene Cluster 5 (n=8 genes: *LDLR, MAPK3, CARD11, GSDMD, SRC, RIPK3, KLF2* and *TICAM1*) and Gene Cluster 6 (n=23 genes: *ACTB, PKM, ACTG1, ACTN4, ENO1, MYH9, CD44, ITGB1, SPP1, TIMP1, ACTN1, AKT2, TKT, F3, AKT1, MFN2, MTOR, MYD88, DNAJB1, CEBPB, DCBLD2, KRT10* and *MAPK7*). Consensus clustering was performed using R v3. 6. 3 package ConsensusClusterPlus, with parameters reps=100, pItem=0. 8, pFeature=1, clusterAlg=”ca”, distance=”euclidean”, seed=123, and maxK=8. For both gene-wise and sample-wise clustering, the optimal clustering solution (each being k=6) was chosen based on the area under the cumulative distribution function (CDF) curve.

### Differential expression and gene set enrichment analysis

Differential expression analysis (DEA) was performed between NET signature group 1 versus group 2 on normalized gene counts using Wilcoxon mean rank-sum tests followed by Benjamini-Hochberg multiple test correction. Gene set enrichment analysis was performed separately on the top 500 most up-and down-regulated DE genes (adjusted p<0. 05) using hypergeometric tests, in which genes were assessed for overlap with each of 32,284 gene sets obtained from the molecular signatures database (MSigDB^69^; downloaded April 2021). Hypergeometric test p values were subjected to Benjamini-Hochberg multiple test correction. Pan-cancer gene-gene relatedness scores (mutual rank) were derived from the GeneFriends dataset^43^ and used to visualize pairwise gene relationships for genes belonging to the Hallmark Epithelial-mesenchymal transition gene set that were up-regulated in the NET signature-high/ITGB1-high group.

### Production and isolation of Neutrophil Extracellular Traps (NETs)

Neutrophil extracellular traps (NETs) were isolated from HL-60 cells (ATCC CCL-240), a peripheral blood pro-myeoloblast that can differentiate into mature neutrophils^40^. Care was taken to ensure cells were sub-cultured at densities that did not induce spontaneous differentiation. For differentiation of HL-60 cells into neutrophils, cells were resuspended in DMSO-containing differentiation medium (phenol-red free RPMI 1640, 10% FBS, 1x glutamax, 1. 5% DMSO) and plated in 10 cm tissue culture-treated plates at a concentration of 2 x 10^5^ cells/mL and incubated for 7 days. Differentiated cells were then activated to undergo NETosis using phorbol 12-myristate 13 acetate (PMA). Cells suspensions from 5 plates were pooled and PMA was added to a final concentration of 1 µM. Cells were mixed by gentle pipetting, re-plated in 10 cm plates and incubated for 4 hours to trigger NETosis. To harvest the NETs, supernatants containing non-adherent cells were removed by aspiration, 1 mL/plate serum-free media was added and NETs were collected by scraping. NETs were pooled, centrifuged at 1000 rpm for 5 minutes to remove cell debris and NET-containing supernatant was transferred to fresh tube. NET-DNA was quantified using a Nanodrop spectrophotometer and stock preparations were dilution to 100 µg/mL for subsequent use in assays. NETs were used fresh for downstream assays.

### Antibodies

The antibodies used in this study are presented below.

*Western Blot and Immunoprecipitation*
| Antibody | Source | Cat # | RRID | Dilution |
| --- | --- | --- | --- | --- |
| ZEB1 Rabbit mAb (D80D3) | Cell Signaling Technology | 3396 | AB_1904164 | 1:1000 |
| Snail (C15D3) Rabbit mAb | Cell Signaling Technology | 3879 | AB_2255011 | 1:1000 |
| GSK-3B (27C10) Rabbit mAb | Cell Signaling Technology | 9315 | AB_490890 | 1:1000 |
| Phospho-GSK-3B (Ser9) Antibody | Cell Signaling Technology | 9336 | AB_331405 | 1:1000 |
| p44/42 MAPK(Erk1/2)(137F5) Rabbit mAb | Cell Signaling Technology | 4695 | AB_390779 | 1:1000 |
| Phospho-p44/42(Erk1/2) (Thr202/Tyr204) Antibody | Cell Signaling Technology | 9101 | AB_331646 | 1:1000 |
| mTOR (7C10) Rabbit mAb | Cell Signaling Technology | 2983 | AB_2105622 | 1:1000 |
| Phospho-mTOR (Ser2448) Antibody | Cell Signaling Technology | 2971 | AB_330970 | 1:1000 |
| CD44 (E7K2Y) XP Rabbit mAb | Cell Signaling Technology | 37259 | AB_2750879 | 1:1000 |
| Phospho-Akt (Ser473) (D9E) XP Rabbit mAb | Cell Signaling Technology | 4060 | AB_2315049 | 1:1000 |
| Phospho-B-Catenin (Ser33/37/Thr41) Antibody | Cell Signaling Technology | 9561 | AB_331729 | 1:1000 |
| Slug (C19G7) Rabbit mAb | Cell Signaling Technology | 9585 | AB_2239535 | 1:500 |
| Vimentin (D21H3) XP Rabbit mAb | Cell Signaling Technology | 5741 | AB_10695459 | 1:5000 |
| ILK1(4G9) rabbit mAb | Cell Signaling Technology | 3856 | AB_2233861 | 1:1000 |
| Integrin B1 (D2E5) rabbit mAb | Cell Signaling Technology | 9699 | AB_11178800 | 1:1000 |
| Mouse Anti-ILK Clone 3/ILK (RUO) | BD Biosciences | 611803 | AB_399283 | 1:1000 |
| Mouse Anti-N-Cadherin | BD Biosciences | 610920 | AB_2077527 | 1:1000 |
| Mouse anti-Akt | BD Biosciences | 610861 | AB_398180 | 1:1000 |
| Mouse Anti-B-Catenin | BD Biosciences | 610154 | AB_397555 | 1:1000 |
| Anti Vinculin Ab, clone VIIF9 (7F9) | Millipore Sigma | MAB3574 | AB_2304338 | 1:2000 |
| Rabbit anti human CCDC25 | Proteintech | 21209-1-AP | AB_10732690 | 1:1000 |
| Goat anti rabbit IgG(H+L) HRP | Jackson | 111-035-003 | AB_2313567 | 1:10,000 |
| anti mouse IgG HRP | Cell Signaling Technology | 7076 | AB_330924 | 1:10,000 |
| CCDC25 Antibody (F-7) AC | Santa Cruz Biotechnology | sc-515201 AC |  | 10ug/IP |
| Integrin B1/ITGB1 (102DF5) | Santa Cruz Biotechnology | sc-73610 | AB_1123630 | 10ug/IP |
| Normal mouse IgG-AC | Santa Cruz Biotechnology | sc-2343 | AB_737178 | 20uL/IP |

*Rac1 Activity Assay*
| Antibody | Source | Cat # | RRID | Dilution |
| --- | --- | --- | --- | --- |
| Anti Rac1 mAb | Cytoskeleton | ARC03 | AB_10709099 | 1:500 |
| Anti Cdc42 mAb | Cytoskeleton | ACD03 | AB_10716593 | 1:500 |

*ITGB1 function blocking Abs*
| **Antibody** | **Source** | **Cat #** | **RRID** | **Dilution** |
| --- | --- | --- | --- | --- |
| Anti Integrin B1 Ab, clone 6S6 | Millipore Sigma | MAB2253 | AB_11210276 | 5-20ug/mL |
| Hamster Anti mouse CD29 | BD Biosciences | 562219 | AB_1115404 | 5-20ug/mL |

*Immunofluorescence*
| **Antibody** | **Source** | **Cat #** | **RRID** | **Dilution** |
| --- | --- | --- | --- | --- |
| Rabbit anti human CCDC25 | Proteintech | 21209-1-AP | AB_10732690 | 1:50 |
| Mouse anti human ILK | Invitrogen | MA5-17099 | AB_2538570 | 1:100 |
| Goat anti human MPO | R&D Systems | AF3667 | AB_2250866 | 1:40 |
| Mouse anti human Paxilin | BD Biosciences | 610051 | AB_397463 | 1:800 |
| Rabbit anti human Cofilin | Cell Signaling | PA5-27627 | AB_2545103 | 1:800 |
| Rabbit anti human ITGB1 | Cell Signaling | D2E5 |  | 1:50/1:100 |
| Rabbit anti human Cit-H3 | Abcam | Ab5103 | AB_304752 | 1:100 |
| Rabbit anti human ILK | Abcam | Ab106955 | AB_11129360 | 1:100 |

### RNA interference

Qiagen FlexiTube gene solution set of 4 siRNAs for ILK were screened to identify efficacious siRNAs to use in our studies. siRNAs were transfected using SilentFect™ (BioRad, 1703360) as per the manufacturer’s instructions. Protein levels were routinely monitored by Western blotting.

### shRNA transduction

The dox inducible knockdown cell lines for ILK and CCDC25 were generated using SMART vector lentiviral dox inducible shRNA plasmids from Horizon Discovery.

HEK293T cells (RRID:CVCL_0045) were seeded at 20,000/cm2. The following day, cells were transfected with 0. 9 μg pSPAX; 0. 1 μg of pVSVG; and 1 μg of the target shRNA plasmid using TransIT-LT1 transfection reagent (Mirus). After 48h, the virus was harvested by passing the media through a 0. 45 μm filter and then concentrating using 5x Lenti Concentrator (Origene). The virus was added to MIA PaCa-2 Luc+ cells or KPCY Luc+ cells along with 10ug/mL polybrene (Sigma) for 72 hours. Antibiotic selection with 2 μg/mL puromycin and 1ug/mL blasticidin (Luc+) for the knockdown cell lines was initiated and cells were continued to be cultured in the presence of selection antibiotics. To induce the expression of the shRNA for each experiment, transfected cells were treated with 1ug/mL doxycycline for 72 hr, refreshed daily, then lysed in RIPA and analyzed by Western blotting.

### *In vivo* bioluminescence imaging

Luciferin was dissolved in saline solution at 15 mg/mL and 10 µL/g body weight was injected intraperitoneally. Ten minutes post-injection, bioluminescence was imaged using an IVIS optical imaging system. Images were normalized and analyzed via Living Image software and signal is presented as total flux in radians (photons/sec/cm2/steradian) over standardized regions of interest.

### Western blotting

Samples were lysed in RIPA buffer (50 mM Tris-HCl, pH 7. 6, 150 mM NaCl, 0. 1% (w/v) SDS) containing 1 mM Na_3_VO_4_, 2 mM NaF and cOmplete protease inhibitor. About 5-10 μg of protein was analyzed by 4%–12% Bis-Tris gradient gel in 1x NuPAGE MOPS SDS running buffer. Gels were transferred to PVDF membranes for 1 hour at 100 V in Tris-Glycine transfer buffer containing 15% methanol. The PVDF membranes were methanol-fixed briefly and incubated with primary antibodies in 2% BSA in TBST overnight at 4°C, followed by washes with TBST for 3×10min and incubation with HRP-conjugated secondary antibody in 2% skim milk in TBST for 1 hour at room temperature. After subsequent washes with TBST, detection was performed by incubation with Supersignal™ West Femto chemiluminescence reagents and visualization using a Chemidoc XRS+ imaging system (Bio-Rad Laboratories). ß-actin or vinculin were used as loading controls. The relative intensities of protein bands were quantified using Image Lab 5. 2. 1 (BioRad Laboratories) and normalized the loading control.

### Co-immunoprecipitation

Cells were washed with 1X PBS and lysed with ice-cold 0. 3% CHAPS buffer (containing 1 mM Na_3_VO_4_, 2 mM NaF and cOmplete protease inhibitor) using a cell scraper followed by centrifugation to remove cell debris. Two milligrams of protein was immunoprecipitated at 4C overnight using either 10 ug of B1INT antibody (Cell Signaling), 20 uL of CCDC25 antibody/agarose conjugate or protein A/G PLUS agarose alone (Santa Cruz Biotechnology). Pellets were washed 5 times with CHAPS buffer, re-suspended in sample buffer, boiled at 100C for 10 minutes before loading on SDS– PAGE for Western blotting as described above.

### Rac1 Activation Assay

Active GTP-Rac1 levels were analyzed using the Rac1 Pull-Down Activation Assay (Cytoskeleton) with 1 mg of lysate as per manufacturer’s instructions. Briefly, cells were seeded in 15cm2 plates at 20,000/cm2 in media containing 10% FBS and allowed to grow for 72 hr to achieve 30% confluence. Cells were then serum starved with 1% FBS for 24 hours, followed by a further 24 hours at 0% FBS before treatment with HL-60 derived NET-DNA for 20 hours, followed by lysis and processing.

### Migration and invasion assay

The scratch wound migration and invasion assays were performed using the Incucyte^TM^ SX3 live cell analysis instrument (Sartorius Biosciences). For invasion assays, 96-well Imagelock® plates (RRID) were coated with a thin layer of phenol red free, growth factor reduced Matrigel (Corning 356231, 50 µL/well of a 100 µg/mL stock solution/well) and incubated for 2 hours at 37°C to allow for matrix polymerization. Following liquid aspiration, MIA PaCa-2 cells or KPCY C1 cells were seeded at a density of 3×10^4^ cells/well in 100 µL/well media and allowed to attach overnight. The next day, confluent cell monolayers were treated with 5 µg/mL mitomycin C (M4287, Millipore Sigma) for 2 hours at 37°C to inhibit cell proliferation. Monolayers were then wounded in a uniform manner with a 96-pin Incucyte Woundmaker™ tool, followed by two successive media changes with 100 µL/well culture media to remove non-adherent cells. Each plate was cooled and the cell monolayer was carefully overlaid with 50 µL/well of 8 mg/mL Matrigel containing 10 µg/mL NETs or equivalent volume of media as a control. The plates were incubated at 37°C for 30 min to polymerize the Matrigel. Finally, 100 µL/well of media containing 10 µg/mL NETs was added on top of the polymerized Matrigel layer. Plates were placed in the Incucyte® Live-Cell Analysis System and images were acquired every 2 h for the indicated times using a Nikon Plan Fluor 10x/0. 3 NA objective in phase contrast mode. Cell invasion was quantified using the Incucyte Scratch Wound Analysis Software Module and data are expressed as relative wound density (%). For samples containing NETs exposed to DNase1, NETs were pre-incubated with 40 U DNase1 per 1 ug DNA for 15 min at 37°C prior to addition to Matrigel layer. Additionally, 1000 U DNase1 in a total media volume of 100 µL/well was added on top of the Matrigel layer. For studies involving pharmacologic inhibition of ILK, QLT-0267 was added at a final concentration of 10 µM in 100 µL/well of media as the top layer. For studies involving incubation with integrin β1-blocking antibodies, the antibody was added at a final concentration of 1 to 5 µg/mL. To account for the 50 µL volume of the Matrigel layer, DNase1, QLT-0267 and integrin β1-blocking antibody were added at 1. 5x the final concentration in 100 µL/well of media. In studies using Dox inducible shRNA stable cell lines, 1 µg/mL doxycycline was added for 72 hours, refreshed daily, to induce shRNA-mediated suppression of gene expression prior to initiation of downstream assays.

### Spheroid growth assay

MIA PaCa-2 cells were seeded at 2,000 cells/well in an ultra-low attachment U bottom 96-well plate (Corning 7007) in 100 µL/well complete media, centrifuged at 800 rpm for 10 min, placed in the Incucyte^TM^ SX3 live cell analysis instrument using the “single spheroid” mode and images were acquired every 6 hours for 9 days. After 72 hours, the newly formed spheroids (approximately 200 μm), were treated with either of 10 µL of media containing 100 µg/mL NETs, 5 µL of 200 μM QLT-0267 in 10% DMSO, or 1000 U of DNase1.

### Preparation of matrix-coated microscope slides

12 mm glass coverslips were coated with Matrigel in DMEM (1:40, Cat no. 356231, Corning), 2 μg/ml of Fibronectin (Cat no. F1056-1MG, Sigma), 0. 01% of Poly L Lysine (Cat no. P47-7, Sigma) or left uncoated, incubated for 1 h at 37°C and washed extensively with PBS.

### Immunofluorescence staining

MIA PaCa-2 dox-inducible shILK cells were cultured for 3 days with or without doxycycline (Cat no. D9891-25G) to induce knockdown of ILK expression. Cells were then harvested, counted, seeded onto coverslips at 7×10^4^ cells/mL and incubated for 6h to allow the cells to adhere to the coverslips. NETs at a concentration of 20 μg/mL and cells were incubated for 24hrs. Confluency checked next day (30-50%). Media was aspirated and cells were washed with Hanks buffered saline (Cat no. 14175-095, Gibco). 4% PFA (Formaldehyde Cat no. YA357388, Thermo) diluted in PBS was added to each well and left at room temperature for 15 minutes. The coverslips in the wells were washed three times with 1X PBS and then blocked using Hanks buffered saline + 10% FBS for 30 min. Coverslips were washed 3 times with 1X PBS. Coverslips were permeabilized with 0. 1% Triton X-100 (Cat no. SLB 56421, Thermo) for 10 min. The coverslips were washed three times with PBS and blocked again using Hanks buffered saline + 10% FBS for 30 mins and washed 3 times with 1X PBS. Coverslips were incubated with X primary antibodies for 24 hours in 4C. AlexaFluor 488/594 conjugated secondary antibodies (1:100, Cat no. A11029/A11012, Life technologies) for 1 h at room temperature (RT). For Dapi staining, the permeabilized cells were incubated with Hoescht (1:10,000, Cat no. 62249) for 10 min and washed with PBS. The coverslips were then mounted in Prolong Diamond anti-fade mounting media (Cat no. P36970, Thermo). Images of cells were acquired using a Zeiss LSM 800 Airyscan confocal microscope (Carl Zeiss, Thornwood, NY) equipped with a 63x oil immersion objective lens and processed using Zen 3. 10 software (Zeiss).

### Tissue immunofluorescence staining

Five micron sections of formalin-fixed, paraffin-embedded tissue were deparaffinized, rehydrated and antigen retrieval was performed by microwaving in 10 mM citrate buffer, pH 6. 0 for 10 minutes. Staining of tissue sections was performed using conditions similar to those used for IF staining of cells. Images of tissue sections were acquired using a Zeiss Colibri inverted microscope (Carl Zeiss, Thornwood, NY) equipped with a 20x lens and processed using Zen 3. 10 software (Zeiss).

### Statistical Analysis

For clinical datasets, One-way ANOVA tests were used to test for differences in a continuous variable across discrete groupings and were followed by Wilcoxon mean rank-sum tests for post-hoc identification of differences between group pairings. Log-rank tests were used to calculate p values in Kaplan-Meier analysis. All comparison tests were two-tailed. All p values were subjected to Benjamini-Hochberg multiple test correction when applicable. Bioinformatics analyses of clinical sequencing data were performed using R v3. 6. 3. For analysis of proteomic data, ANOVAs were carried out on specific proteins using a previously published proteomic data set^38^ and methods described previously^38^.

For analysis of non-clinical datasets, the data were presented as Mean ± SD or as Mean ± SEM when multiple experiments are represented. Statistical analyses were conducted using Prism software (GraphPad, La Jolla, CA). Statistical significance for multiple comparisons was calculated using two-way ANOVA corrected for multiple comparisons with the Holm-Sidak method unless otherwise indicated. Student’s t-tests corrected for multiple comparisons using the Holm-Sidak method was used when only two data sets were being compared in a given plot. For all tests, the significance is indicated as *p < 0. 05; **p < 0. 01; ***p < 0. 001.

### Materials availability

All requests for resources and reagents should be directed to and will be fulfilled by the lead contact as designated above. This includes antibodies and shRNA engineered cell lines. Genetically engineered cells lines, the PaCa41 PDX cell line and QLT-0267 will be made available on request after completion of a Materials Transfer Agreement. Sequencing data generated as part of the PanGen/POG study is available in the European Genome-phenome Archive (EGA) under accession number #EGAS00001001159.

## Acknowledgements

This work was supported by grants from the Canadian Institutes of Health Research (CIHR; Foundation Scheme Grant FDN-143318 to S. D., Project Grant 486353 to S. D., D. J. R. and D. F. S.) and the Cancer Research Society (CRS; Rob Lutterman Memorial Fund Operating Grant 938669 to S. D.). Funding was also provided by the BC Cancer Foundation (BCCF) to S. D.

We gratefully acknowledge the participation of patients and their families, and the PanGen and POG teams. The results published here are, in part, based upon data generated by the International Cancer Genome Consortium (https://icgc.org/). Data used in this publication were generated in part by the National Cancer Institute Clinical Proteomic Tumor Analysis Consortium (CPTAC). This research was supported with funding from BC Cancer Foundation and Terry Fox Research Institute (Project 1078) (to D. J. R. and D. F. S.).

## Author Contributions

*Conceptualization:* P. C. M., J. TT., and S. D.

*Methodology*: P. C. M., J. T. T., S. A., H. T., R. C., Z. J. G., W. S. B., S. E. K., J. M. K, and S. D.

*Investigation:* P. C. M., J. T. T., S. A., H. T., R. C., Z. J. G., W. S. B., S. E. K., J. M. K.

*Formal analysis:* P. C. M., J. T. T., S. E. K., and J. M. K.

*Data Curation:* P. C. M, J. T. T.

*Resources:* P. T., R. G., S. J. M. J, J. L., M. A. M., G. B. M. D. J. R, D. F. S, and S. D.

*Writing – Original draft:* P. C. M. and S. D.

*Writing – Review & Editing:* P. C. M., J. T. T., S. A., J. M. K., and S. D.

*Supervision*: P. C. M., D. J. R, D. F. S, and S. D.

*Funding acquisition:* D. J. R., D. F. S., and S. D.

---

**Extended Data Fig. 1.**
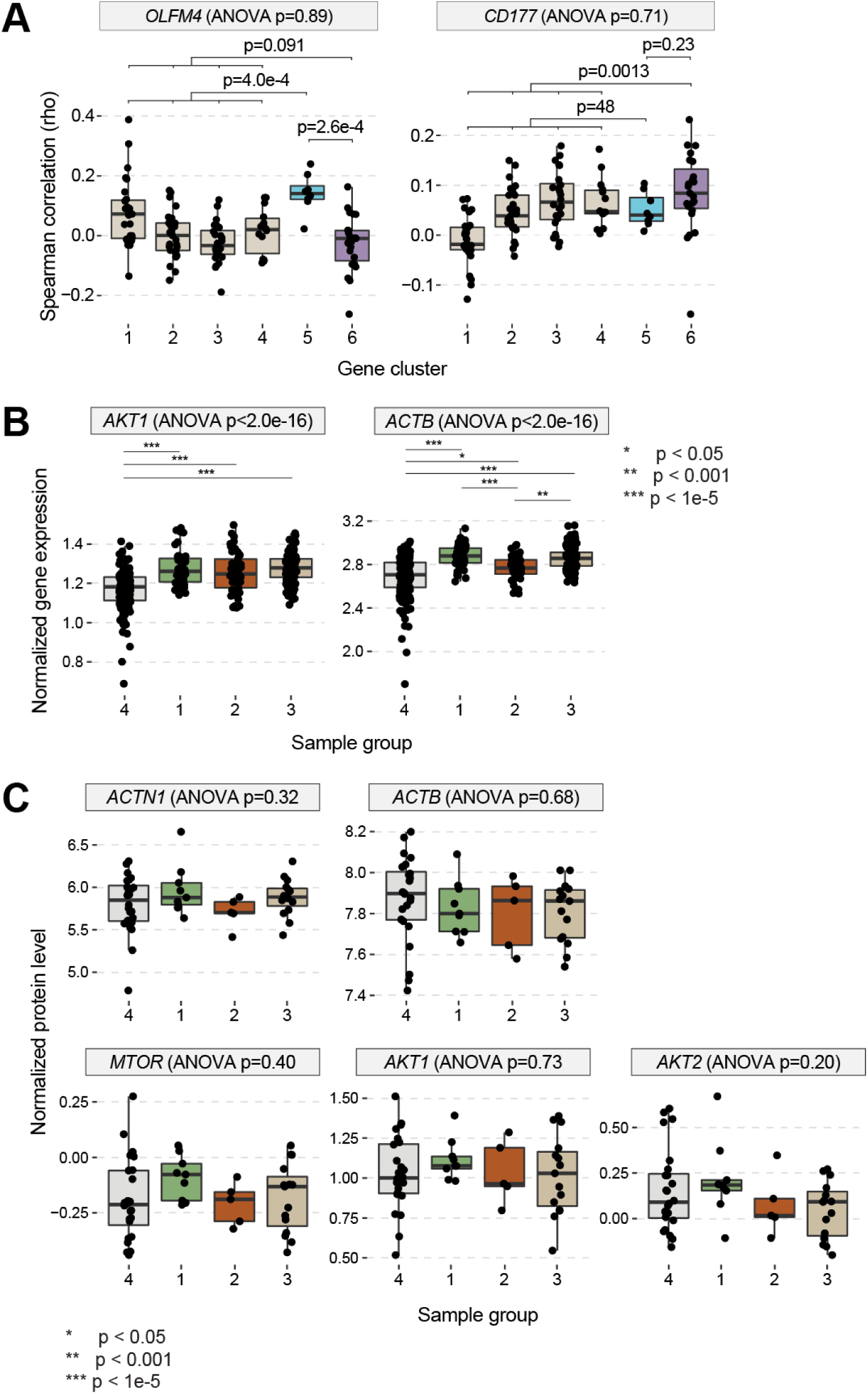
**(A)** Box plots showing the relationship between expression of the indicated genes and each gene cluster identified in Figure 1B. **(B, C)** Box plots showing (B) normalized gene expression levels and (C) normalized protein levels for each of the indicated genes across patient groups 1 to4 identified in Figure 1D.

**Extended Data Fig. 2.**
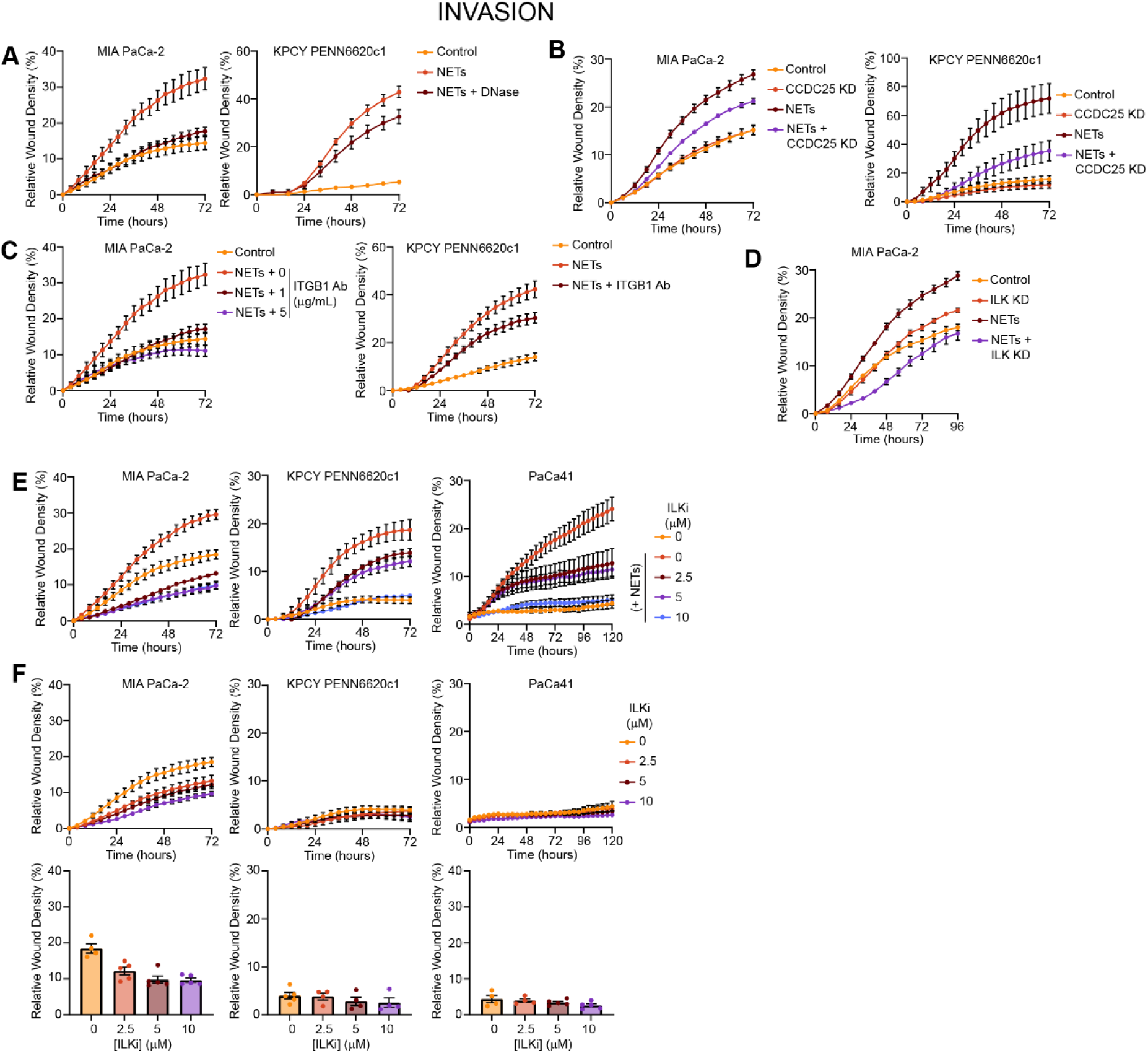
**(A)** Kinetic curves showing invasion through matrigel over time of the indicated cell lines exposed to NETs incubated with and without DNase. **(B)** Kinetic curves showing NET-induced invasion through matrigel over time by human and mouse PDAC cells following knockdown of CCDC25 expression using dox-inducible shRNA. **(C)** Kinetic curves showing NET-induced invasion through matrigel over time by the indicated cell lines cultured with or without function-blocking antibodies targeting ITGB1. **(D)** Kinetic curves showing NET-induced invasion through matrigel of MIA PaCa-2 cells following depletion of ILK expression using siRNA. **(E)** Kinetic curves showing invasion through matrigel of the indicated cell lines in the presence of a specific inhibitor of ILK, QLT-0267. (F) Invasion through Matrigel by the indicated cell lines in basal conditions in the presence or absence of the ILK inhibitor. Top plots show kinetic curves for invasion over time. Bottom plots show levels of invasion at the 72 hour time point.

**Extended Data Fig. 3.**
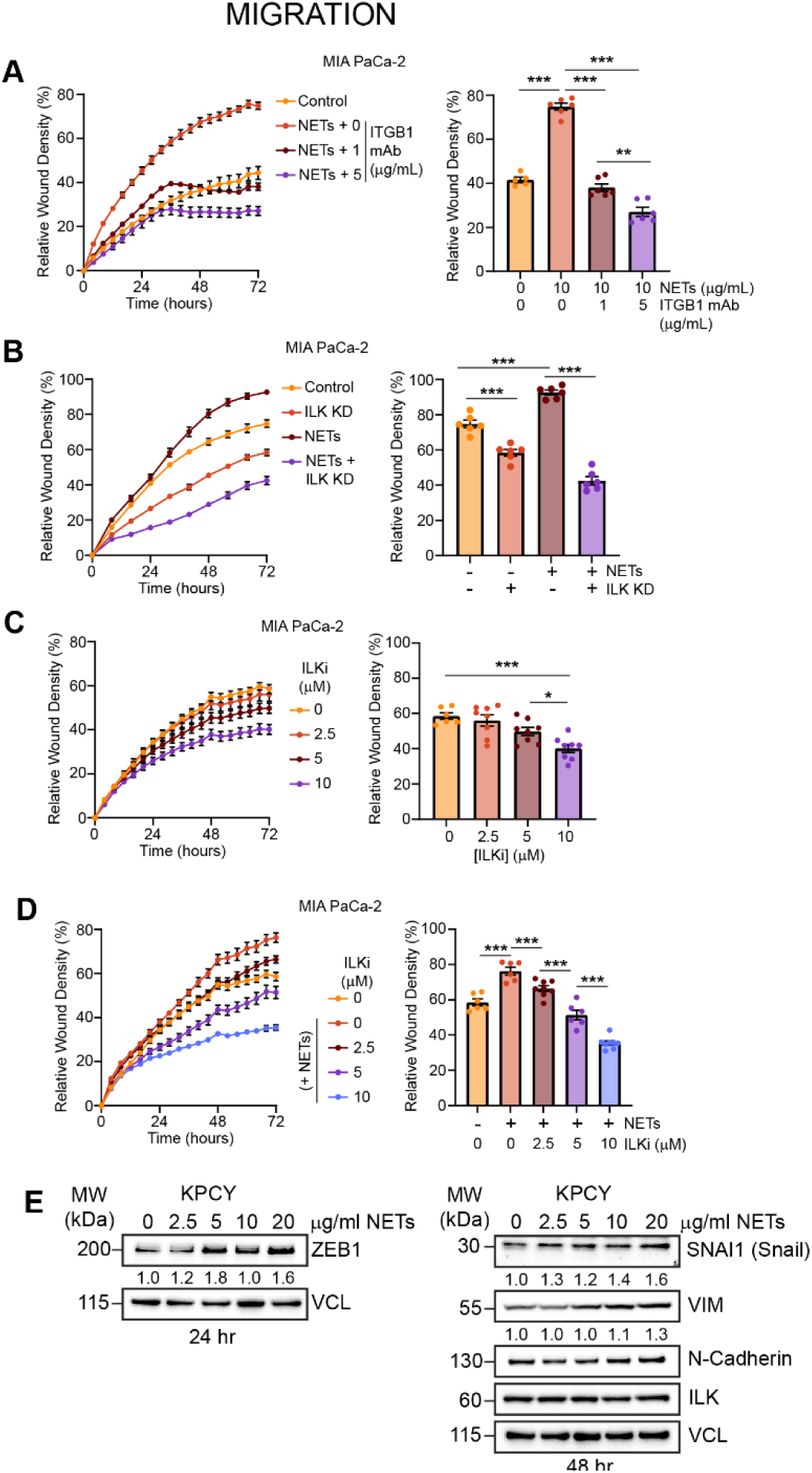
**(A)** Graphs showing NET-induced migration of MIA PaCa-2 cells cultured with or without function-blocking antibodies targeting ITGB1. **(B)** Graphs showing migration of MIA PaCa-2 cells following depletion of ILK expression using siRNA. **(C)** Graphs showing baseline levels of migration of MIA PaCa-2 cells cultured in the presence of increasing concentrations of the specific ILK inhibitor, QLT-0267. **(D)** Graphs showing migration of NET-induced migration of MIA PaCa-2 cells cultured with increasing concentrations of the ILK inhibitor. **(E)** Western blot analysis of the indicated EMT markers in KPCY cells in the presence of increasing concentration of NETs at the indicated time points. For all panels, the kinetic curves are shown to the left and the level of migration at 72 hours is shown to the right. \**P* < 0. 05; \*\**P* < 0. 01; \*\*\**P* < 0. 001.

**Extended Data Fig. 4.**
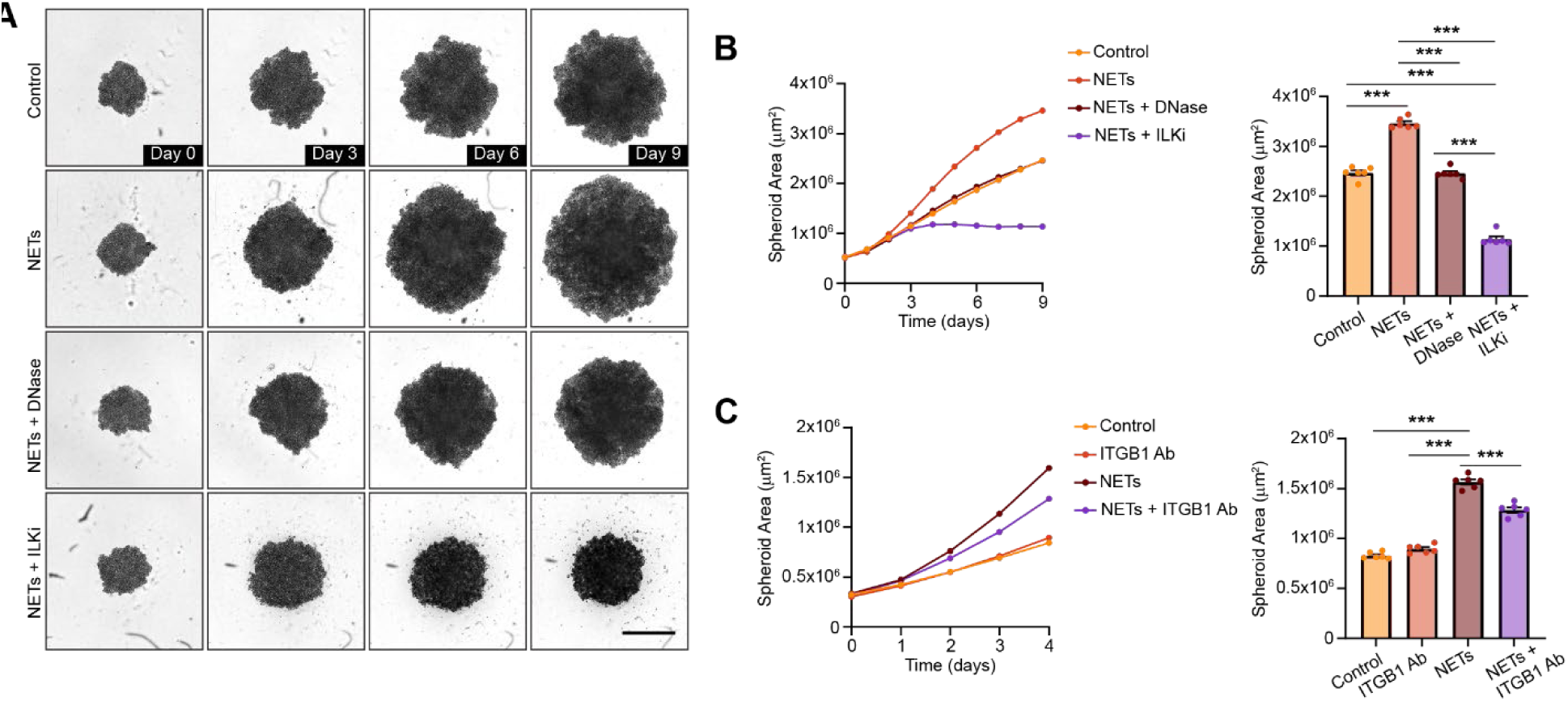
**(A)** Time lapse images of MIA PaCa-2 spheroids cultured with or without NETs treated with DNase I or with NETs in the presence of the ILK inhibitor. **(B)** Quantification of spheroid area, a metric of spheroid size, for the spheroids cultured as described in panel A. The kinetic curves are shown to the left and the analysis of spheroid area at day 9 is shown to the right. **(C)** Graphs showing the change in size over time of MIA PaCa-2 spheroids cultured with or without NETs in the presence or absence of function-blocking antibodies targeting ITGB1. \*\*\**P* < 0. 001.

**Extended Data Table 1.** Set of NET-related genes used for gene-wise co-expression cluster analysis of human PDAC samples

| Set of NET-related genes |  |  |  |  |  |
| --- | --- | --- | --- | --- | --- |
| ACTB | CLEC6A | GPBAR1 | LDLR | NLRP3 | SOCS3 |
| ACTG1 | CLEC7A | GSDMD | LPAR3 | NOX4 | SPP1 |
| ACTN1 | CSF3 | HIF1A | LTF | OPA1 | SRC |
| ACTN4 | CTSC | HMGB1 | LYZ | P2RX1 | STAT3 |
| AKT1 | CTSG | HRG | MAPK1 | PADI4 | SUCNR1 |
| AKT2 | CXCL1 | IL12A | MAPK14 | PARVB | SYK |
| ATG7 | CXCL2 | IL17A | MAPK3 | PF4 | TICAM1 |
| AZU1 | CXCL5 | IL1B | MAPK7 | PIK3CA | TIMP1 |
| C3 | CXCR4 | IL33 | MCOLN3 | PKM | TKT |
| C3AR1 | CYP27A1 | IL5 | MFN1 | PRKG2 | TLR2 |
| C5AR1 | DCBLD2 | IL6 | MFN2 | PROCR | TLR7 |
| CAMP | DNAJB1 | IL8 | MMP3 | PRTN3 | TLR8 |
| CARD11 | DNASE1 | IRAK4 | MMP9 | PTAFR | TNF |
| CASP1 | ELANE | IRF1 | MNDA | RIPK1 | TNFAIP3 |
| CAT | ENO1 | ITGAM | MPO | RIPK3 | VCAM1 |
| CCL2 | ENTPD4 | ITGB1 | MTOR | S100A12 | VENTX |
| CD274 | F2RL2 | ITGB2 | MYD88 | S100A8 | WASL |
| CD44 | F3 | KCNN3 | MYH9 | S100A9 | XIST |
| CEBPB | FCAR | KLF2 | NFE2L2 | SELP |  |
| CFTR | FCGR2B | KRT10 | NFIL3 | SELPLG |  |
| CLEC4E | FGL2 | LCP1 | NFKBIA | SGK1 |  |

